# The conserved HIV-1 spacer peptide 2 triggers matrix lattice maturation

**DOI:** 10.1101/2024.11.06.622200

**Authors:** James C. V. Stacey, Dominik Hrebík, Elizabeth Nand, Snehith Dyavari Shetty, Kun Qu, Marius Boicu, Maria Anders-Össwein, Robert A. Dick, Walther Mothes, Hans-Georg Kräusslich, Barbara Müller, John A. G. Briggs

**Affiliations:** Department of Cell and Virus Structure, Max Planck Institute of Biochemistry, 82152 Martinsried, Germany; Structural Studies Division, MRC Laboratory of Molecular Biology, Cambridge CB2 0QU, United Kingdom; Department of Microbial Pathogenesis, Yale University School of Medicine, New Haven, CT 06536, USA; Department of Infectious Diseases, Virology, Heidelberg University, 69120 Heidelberg, Germany; Infectious Diseases Translational Research Programme, Department of Biochemistry, Yong Loo Lin School of Medicine, National University of Singapore, 117545, Singapore; Department of Molecular Biology and Genetics, Cornell University, Ithaca, NY 14853, USA; German Center for Infection Research, Heidelberg, Germany

**Author notes:** Department of Pediatrics, School of Medicine, Emory University, Atlanta, GA, USA. These authors contributed equally to this work (James C. V. Stacey, Dominik Hrebík). Correspondence to JAGB.

## Abstract

HIV-1 particles are released in an immature, non-infectious form. Proteolytic cleavage of the main structural polyprotein Gag into functional domains induces rearrangement into mature, infectious virions. In immature virus particles, the Gag membrane binding domain, MA, forms a hexameric protein lattice that undergoes structural transition upon cleavage into a distinct, mature MA lattice. The mechanism of MA lattice maturation is unknown. Here we show that released spacer peptide 2 (SP2), a conserved peptide of unknown function situated ∼300 residues downstream of MA, binds MA to induce structural maturation. By high-resolution in-virus structure determination of MA, we show that MA does not bind lipid into a side pocket as previously thought, but instead binds SP2 as an integral part of the protein-protein interfaces that stabilise the mature lattice. Analysis of Gag cleavage site mutants showed that SP2 release is required for MA maturation, and we demonstrate that SP2 is sufficient to induce maturation of purified MA on lipid layers in vitro. SP2-triggered MA maturation correlated with faster fusion of virus with target cells. Our results reveal a new, unexpected interaction between two HIV-1 components, provide a high-resolution structure of mature MA, establish the trigger of MA structural maturation, and assign function to the SP2 peptide.

## Introduction

HIV-1 morphogenesis proceeds via assembly of the viral polyprotein Gag at the plasma membrane of the infected cell. Gag-Gag, Gag-membrane, Gag-RNA and Gag-host protein interactions drive bud formation and release of an immature, non-infectious viral particle from the cell surface. Concomitant with budding, Gag undergoes an ordered proteolytic cleavage cascade mediated by the viral protease (PR) into its subdomains MA (matrix), CA (capsid), spacer peptide 1 (SP1), NC (nucleocapsid), spacer peptide 2 (SP2) and p6 (Extended Data Fig. 1a). This leads to structural changes in the protein domains and a dramatic rearrangement of viral architecture ^1-4^. Maturation converts a non-infectious particle optimised for assembly into a virion specialised and competent for entry and infection ^1-3,5^. The maturation of the CA lattice has been studied in detail providing models for how cleavages upstream and downstream of CA trigger disassembly of the spherical, hexameric immature CA lattice and reassembly into a structurally and functionally distinct mature CA lattice ^3,4,6-9^. These analyses established a key role for SP1 release in CA lattice maturation.

The MA domain of Gag is responsible for the recruitment of Gag to the host plasma membrane during virus assembly ^10-13^. MA forms a trimer that interacts with membranes in a phosphatidylinositol 4-5 bisphosphate (PI(4,5)P_2_)-dependent manner via a highly basic region (HBR) and an N-terminal myristate moiety ^14,15^. Myristate is initially sequestered within MA, but is exposed upon MA trimerisation, and can insert into the inner leaflet of the bilayer ^14-16^ (Extended Data Fig. 1b). Mutations in MA lead to defects in envelope protein (Env) incorporation, but the mechanisms by which MA contributes to Env incorporation are not fully resolved ^10,17-21^. We recently showed that trimer-trimer interactions between N-terminal residues link MA trimers together into a loose hexameric lattice in the immature virion ^22^. Upon maturation, these interactions are replaced by a larger trimer-trimer interface between MA trimers to form a distinct, regularly packed hexameric lattice in the mature virion ^22^. In contrast to CA, high-resolution structural data on the immature and mature MA lattices are not available.

Nuclear magnetic resonance (NMR) studies of monomeric MA previously revealed that a cleft in MA formed by α-helix_4_ and the loop between α-helix_1_ and α-helix_2_ can bind the inositol head group and one of the acyl chains of PI(4,5)P_2_. On a membrane, this binding mode would require partial extraction of PI(4,5)P_2_ from the bilayer (Extended Data Fig. 1b). In vitro, PI(4,5)P_2_ binding promotes exposure of the MA-myristoyl moiety ^11^. Cryo-electron tomography of intact virions showed that this cleft is empty and exposed in the immature virus but occupied and facing the trimer-trimer interface in the mature MA lattice ^22^. The cryo-electron tomography density observed in the cleft in the mature virus was consistent with the bound PI(4,5)P_2_ observed by NMR, but the low resolution of the tomography structure prevented a conclusive assignment ^22^. The same cleft has also been implicated in the binding of host tRNAs ^23^. Bound tRNA protrudes above the membrane binding surface of MA, resulting in the model that the MA domain of cytosolic Gag binds tRNA which is displaced by membrane binding during bud assembly ^23^ (Extended Data Fig. 1b). Displacement would allow the membrane binding surface to interact with PI(4,5)P_2_ and other lipids in the bilayer, and free the cleft for binding to (partially membrane extracted) PI(4,5)P_2_ during the later maturation step ^22^. Taken together with the observation that both the mechanical properties of the viral membrane ^24,25^ and virion fusogenicity ^5,26^ change upon maturation, these results led to the hypothesis that MA maturation changes the properties of the viral envelope bilayer by extracting lipids.

While these results determined structural features governing Gag trafficking and membrane binding, the mechanism of structural maturation of the MA lattice in the released virion remains unknown. The simple assumption would be that MA maturation is induced when it is released as a mature protein domain from Gag by cleavage between MA and CA. However, we previously observed that this is not the case: a Gag mutant in which cleavages between MA and CA and between CA and SP1 are blocked (MA-SP1) displays an immature CA lattice and a mature MA lattice, suggesting that long range interactions may be important for MA lattice maturation.

Here we set out to determine the mechanism by which MA maturation is induced, to identify how lipid binding is mediated and changed during maturation, to define the interactions that mediate immature and mature MA lattice formation, and to investigate whether these phenotypes influence the fusion properties of HIV-1. We have obtained high-resolution structures of the MA lattice within intact wild-type (WT) virus particles and have studied Gag cleavage mutants. Unexpectedly, our data show that the previously described mature MA ligand is not a lipid; instead, it is SP2, a highly conserved downstream Gag peptide of unknown function. Proteolytic release and MA binding of SP2 is the trigger for MA maturation and correlates with the virus gaining fast, WT-like fusion kinetics *in vitro*.

## Results

### High-resolution structures of MA within viruses

To determine high resolution structures of the immature and mature MA lattices within intact viruses we applied *in situ* single-particle analysis (SPA). We and others recently used a similar approach to determine the structure of the mature HIV-1 CA lattice *in vitro* at high resolution ^27-29^. SPA is faster than our previous cryo-electron tomography approach ^22^, allowing us to collect and analyse larger datasets and to obtain reconstructions of the MA lattice from immature (PR-) virus particles at 5.8 Å, from mature virus particles at 3.8 Å and from MA-SP1 virus particles (where MA also adopts a mature lattice) at 3.1 Å (Fig. 1b-i, Extended Data Fig. 2-6). The lower resolution of the immature lattice likely reflects its higher flexibility. In all cases, the results were consistent with our previous lower resolution cryo-ET structures at the resolution at which they could be compared ^22^. However, the substantially higher resolution of the structures described here unravelled crucial structural details.

**Fig. 1.**
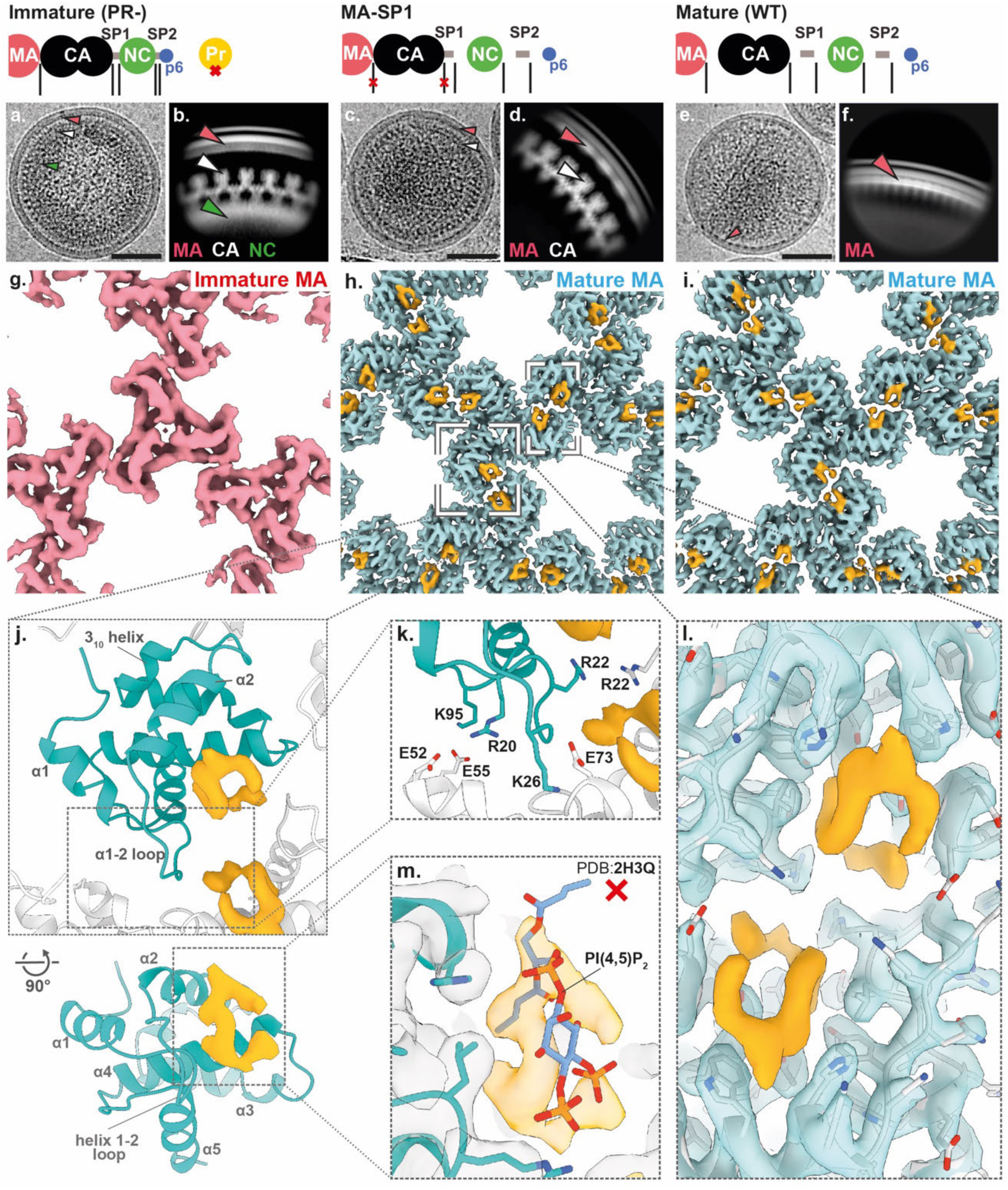
High resolution reconstructions of the Immature, Mature and MA-SP1 MA lattices. **a,** Representative Immature (PR-) HIV-1 virion from single particle cryo-EM data. Arrow heads highlight specific Gag domain layers that are observed (red MA; white CA and green NC) Scale bar, 50nm. **b,** Side-view 2D class of the Gag layer from single particle data of the Immature (PR-) virions. **c,** Representative MA-SP1 HIV-1 virion from single particle data. **d,** Side-view 2D class of the Gag layer from single particle data of MA-SP1 virions. As expected, the cleaved NC layer is not observed in 2D classes. **e,** Representative Mature (WT) HIV-1 virion from single-particle cryo-EM data. **f,** Side-view 2D class of the Gag layer from single particle data of Mature (WT) virions. As expected, only the MA layer is observed on the membrane. For the three datasets (Immature, MA-SP1 and Mature) a schematic representation of the expected Gag cleavage state is provided. **g-i,** Respective 3D cryo-EM reconstructions of the MA lattice determined from data, viewed from the membrane. Consistent with (Qu et al. 2020), the MA lattice was found to be immature in Immature (PR-) particles, mature in MA-SP1 particles and mature in Mature (WT) particles. Additional ligand density is observed (orange) in both mature lattices, but was absent in the immature lattice. **j,** Top view of the model of the mature MA lattice, including the side pocket ligand density, with a single monomer coloured blue and surrounding MA monomers coloured white. Lower panel shows rotated view of MA monomer with helices labelled. **k,** Zoom in view of the trimer-trimer interface in the atomic model of the mature matrix lattice. The interface is mediated by specific electrostatic interactions between HBR residues and an electro-negative patch of residues distributed along the intra-trimeric interface. **l,** Zoomed in view of h showing the ligand binding across the trimer-trimer interface with fit atomic model (grey). **m,** Rigid fitting of the deposited model of MA bound to PI(4,5)P_2_ (PDB: 2H3Q) ^11^, shows that the density is not consistent with PI(4,5)P_2_ binding in the manner previously reported (the PI(4,5)P_2_ atomic model is not accommodated within the orange ligand electron microscopy density).

In the mature MA lattice, the ordered loop between α-helix_1_ and α-helix_2_ sits at the centre of the inter-trimer interaction, forming a 230 Å^2^ interface with, in particular, helix_4_ in its two-fold related neighbour and a 90 Å^2^ interface including the 3_10_ helix of another MA molecule. The loop between α-helix_1_ and α-helix_2_ overlaps with the previously described highly-basic-region (HBR) of MA, and the structure revealed HBR binding into an acidic surface in the adjacent trimer: the positions of residues R20, K26 and K95 allow them to form inter-trimer salt bridges with E52, E73/E74 and E55 respectively (Fig. 1j-k). Sandwiched between the two-fold related neighbours is the ligand-bound side pocket where the two symmetry-related ligand densities come into close contact, bridged by residue R22 (Fig. 1k,l).

The shape of the observed density for the ligand bound to the MA side pocket (Fig. 1m) was in poor agreement with the binding conformation of PI(4,5)P_2_ as previously observed by NMR ^11^. We therefore explored alternative positions of PI(4,5)P_2_, but were unable to obtain a fit for PI(4,5)P_2_ or for other stoichiometrically relevant lipid species consistent with the shape of the observed density (Extended Data Fig. 7).

### Influence of Gag processing on the MA lattice

The observation that the HIV-1 variant MA-SP1 displays a mature MA lattice suggests that it is not the separation of MA from CA, but other proteolytic cleavage(s) in the C-terminal region of Gag between SP1 and p6 that are relevant for MA lattice maturation. To identify processing steps involved in MA lattice maturation, we produced virus particles for four additional cleavage site mutants: MA-SP2, MA-NC, MA-SP1:NC-p6, and NC-p6 (Fig. 2a, Extended Data Fig. 2a), and characterized their MA lattices by cryo-electron microscopy (Fig. 2b, Extended Data Fig. 2b).

**Fig. 2.**
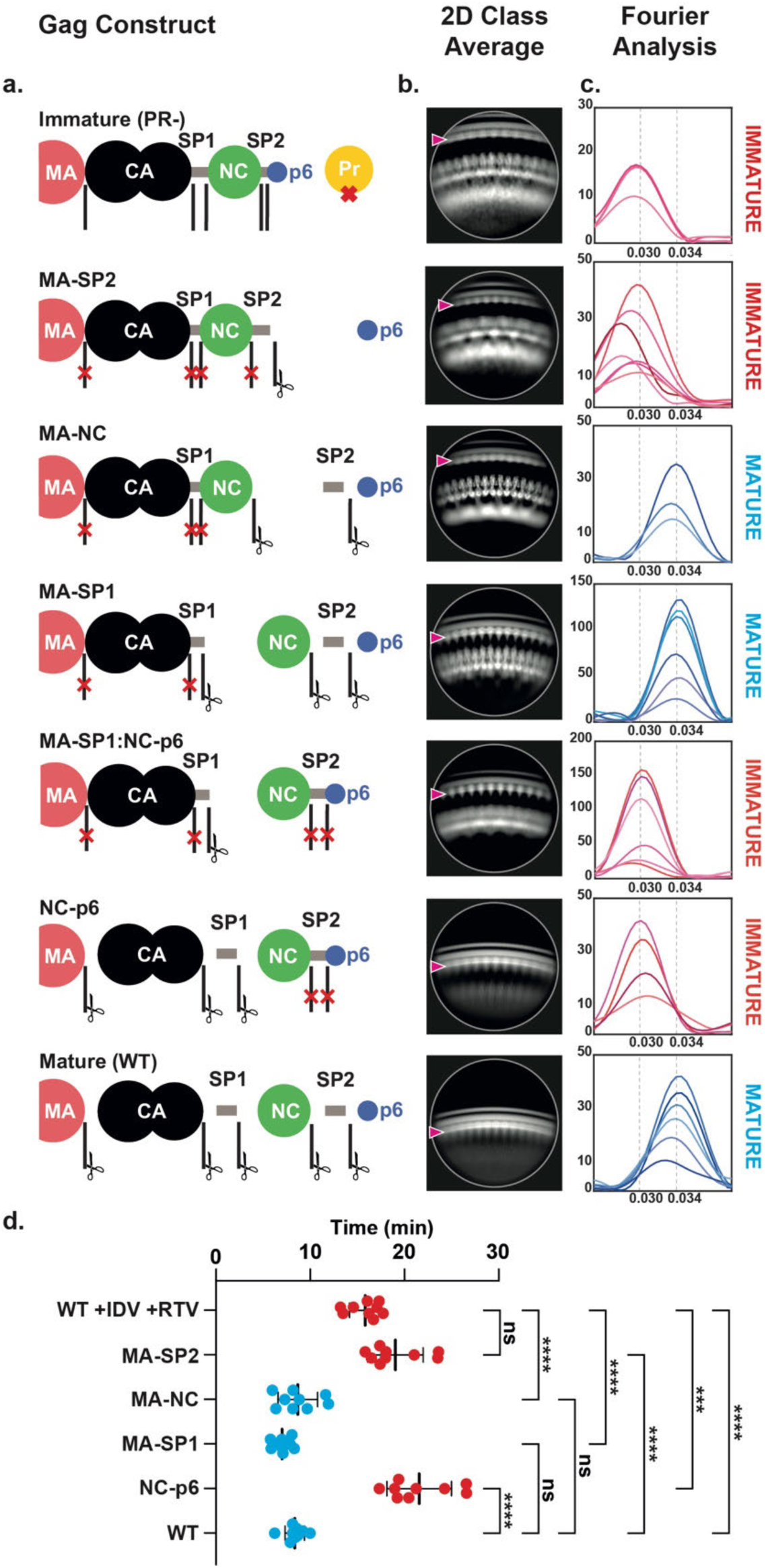
MA lattice states and fusion kinetics for tested Gag cleavage mutants. **a,** Schematic representations of the expected Gag cleavage state in all tested HIV-1 constructs. **b,** Representative side view 2D classes of Gag. Clear ordering is observed within the membrane bound MA lattices (red arrow). The observed MA, CA and NC densities match those expected based on the cleavage state **c,** Fourier analysis of MA lattice spacings from side-views of 2D class averages of HIV cleavage mutants. A Fourier peak with Miller indices of h,k = [2,1] at a spatial frequency of 0.030 A^-1^ represents the lattice spacing of an immature lattice while a Fourier peak with the same Miller indices at a spatial frequency of 0.034 A^-1^ represents the lattice spacing of a mature lattice (Extended Data Fig. 8). Each curve represents a single 2D class average. **d.** Histogram of fusion T_1/2_ for the indicated virus-like particles. All particles contain WT JRFL Env. The first row is immature particles with WT Gag produced in the presence of protease inhibitors Indinavir (IDV) and Ritonavir (RTV). Plotted points are from three biological replicates with three technical replicates per biological replicate for an n = 9. Each biological replicate represents one aliquot from a bulk preparation of viral particles. Statistics were performed using an ordinary one-way ANOVA test. Brown-Forsythe test and Bartlett’s test were performed as corrections. Bars are mean ± s.d.. NS, *p* ≥ 0.10; ***, 0.001 > *p* ≥ 0.0001; ****, *p* < 0.0001. Red and blue points represent cleavage mutants observed in (**c**) as having immature or mature MA, respectively. See also Extended Data Fig. 9.

We analysed 2D class averages of regions of the particle edges that showed clear, repetitive MA densities and measured the repetitive spacing of the MA layer using a Fourier analysis (Fig. 2b,c, Extended Data Fig. 8). This measurement is reference-free and independent of 3D alignment. The approximate hexamer-hexamer distance measured from the immature and mature lattice structures was 10.1 nm and 9.0 nm, respectively. The immature and mature class averages showed a Fourier peak at a spatial frequency of 0.030 Å^-1^ or 0.034 Å^-1^, respectively (Fig. 2c), corresponding to the frequency predicted for the [2,1] lattice reflection of the respective lattices (Extended Data Fig. 8c). MA-SP1, as expected, also showed the 0.034 Å^-1^ peak (Fig. 2c) corresponding to the mature MA lattice. We performed the same analysis on images of the additional cleavage mutants and found that MA-NC has a mature MA lattice while MA-SP1:NC-p6, NC-p6 and MA-SP2 displayed immature MA lattices (Fig. 2c).

The finding that cleavage mutants MA-SP1:NC-p6 and NC-p6 had retained an immature MA lattice (Fig. 2c) indicated that cleavage within the NC-SP2-p6 moiety, rather than separation of this moiety from MA-SP1, is required for MA lattice maturation. The observation that MA-NC had a mature MA lattice structure while MA-SP2 had an immature MA lattice, indicated that release of free SP2 peptide correlates with MA maturation.

### SP2-induced MA maturation correlates with WT-like fusion kinetics

Efficient HIV-1 fusion was previously shown to depend on Gag maturation ^5^. We therefore used the same Gag cleavage mutants and a split nanoluciferase complementation assay to measure HIV-1 fusion. Membrane fusion of immature particles (produced in the presence of protease inhibitors) was two-fold slower than observed for WT mature particles in this assay (Fig. 2d, Extended Data Fig. 9). Using the Gag cleavage mutants, we found that membrane fusion remained slow when SP2 was not released (Fig. 2d). In contrast, fusion was accelerated two-fold to the level of WT particles when SP2 was released from Gag. The release of SP2 and MA maturation thus correlate with WT-like fusion kinetics of HIV-1.

### Free SP2 binds to MA within the virion

The analyses of cleavage mutants raised the question of how release of the distant SP2 peptide may affect maturation of MA bound to the plasma membrane. Speculating that direct binding of SP2 to MA could be the trigger of MA maturation, we asked whether the density previously observed in the side pocket of mature MA ^22^, initially presumed to correspond to a lipid, could instead represent the SP2 peptide. To assess this, we built continuous stretches of amino acids from SP2 into the density. We found that, indeed, the six sequential C-terminal residues (GRPGNF) of SP2 could be confidently fitted into the cryo-EM density in a manner consistent with reconstructions of the mature MA lattice from WT, MA-SP1 and MA-NC particles (Fig. 3a,b, Extended Data Fig. 10a). Secondly, we employed fully automated model building (Relion5: ModelAngelo) with full-length Gag as the input sequence. Since ModelAngelo does not take symmetry into account, we analysed sequences built into each of the three 3-fold symmetry-related densities of the central MA trimer independently. In all three instances, the side pocket density was predicted to correspond to the C-terminal residues of SP2 residues, and the models faithfully recapitulated our initial modelled conformation (Extended Data Fig. 10b,c). Thirdly, we performed the same automated model building without any input sequence, and again in one position obtained the peptide sequence GRPGNF in a conformation closely matching to our final model (Extended Data Fig. 10b,c).

**Fig. 3.**
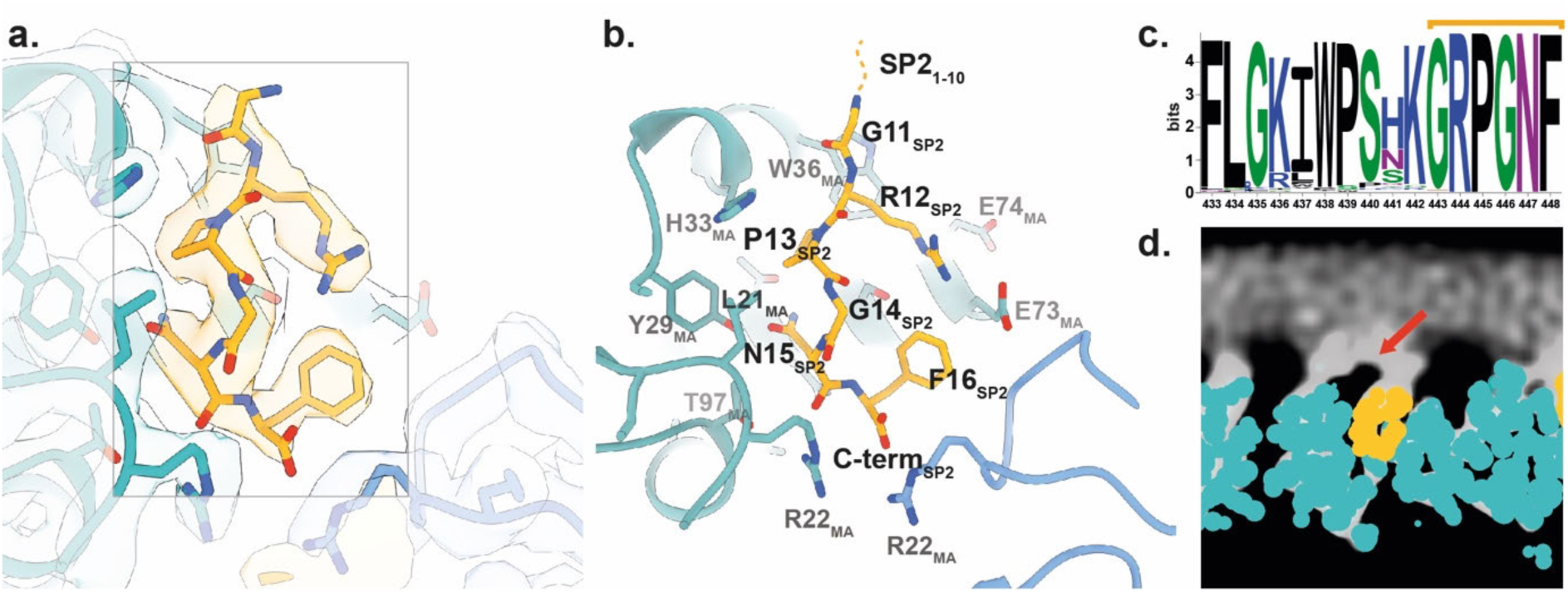
Structure of the SP2 C-terminus bound to the mature MA lattice. **a,** Fit of the six C-terminal SP2 residues (orange) and MA (blue) within the MA-SP1 mature MA lattice reconstruction, showing cryo-EM density. **b,** Same view as A. with residues MA and SP2 residues labelled. **c,** Weblogo representation of the conserved amino acid sequence of SP2, generated using the hiv.lanl.gov filtered web sequence database. Orange bar indicates the six C-terminal residues that are resolved bound to the MA side pocket. **d,** Additional density (red arrow), continuous with that of bound SP2, is observed above MA and contacting the inner leaflet of the viral membrane. Shown as orthoslice of the unsharpened map, with MA and SP2_11-16_ atomic coordinates overlayed as spheres (blue and orange respectively).

In the resulting model, bound SP2 adopted a largely extended conformation, with G11 at the top close to the viral membrane, and the C-terminal F16 at the base (Fig. 3b). The peptide is held in place by multiple interactions, including π-stacking between the backbone amides of SP2 R12 and P13 with MA W36 and H33, respectively, and a salt bridge between SP2 R12 and MA E73. A cleft consisting of residues L21, Y29, S77, T81 and T97 supports the binding of SP2 residues G14 and N15 (Extended Data Fig. 11). Unlike the observed binding mode of tRNA^Lys3^ ^23^ and PI(4,5)P_2_ ^11^, R76 and K27 do not form an electrostatic interaction with SP2, adopting a markedly different conformation (Extended Data Fig. 12). SP2 appears to contribute to formation of the inter-trimer interactions in the mature MA lattice via a salt bridge between the C-terminal carbonyl and R22 in the two-fold related MA molecule (Extended Data Fig. 11c).

The ten N-terminal residues of SP2 were not resolved in our structures. This is not due to proteolytic cleavage within SP2, as the presence of the 16 amino-acid peptide in virions has been demonstrated ^30,31^. The orientation of the peptide would position SP2 residues 1-10 at the viral membrane. Re-examination of the mature MA lattice reconstruction revealed additional diffuse density directly above MA monomers in all three reconstructions (WT, MA-SP1 and MA-NC) (Fig. 3c). This density is tightly associated with the membrane inner leaflet; we hypothesise that it partially represents the N-terminus of SP2 (Fig. 3c,d).

### In vitro reconstitution of MA maturation

The findings described above indicate that the SP2 peptide is necessary to induce MA lattice maturation. We next asked whether SP2 is sufficient to induce formation of a mature MA lattice using an *in vitro* reconstituted system. For this, we added purified, recombinant myristoylated MA protein in the presence or the absence of SP2 to lipid monolayers with a composition mimicking the inner leaflet of an HIV-1 membrane ^32^. Three independently prepared MA-coated lipid monolayers formed on holey-carbon EM grids were imaged by cryo-electron microscopy for each condition. For each grid, thirty images were collected from each of 5 randomly selected grid-squares. A grid of sub-images was extracted from each image resulting in 139,500 sub-images per grid which were subjected to 2D classification. Classes showed either no protein lattice, immature-like MA lattices or mature-like MA lattices (Fig. 4a-c, Extended Data Fig. 13). The total number of sub-images contributing to immature and mature-like MA lattice classes was quantified for three independent experiments (Fig. 4d). In the absence of SP2, 7% of sub-images contributed to immature-like classes while 7% contributed to mature-like classes. The remaining sub-images were assigned to classes containing no interpretable MA lattice. In contrast, 47% of sub-images in the presence of SP2 contributed to mature-like MA classes, while no immature-like classes were observed.

**Fig. 4.**
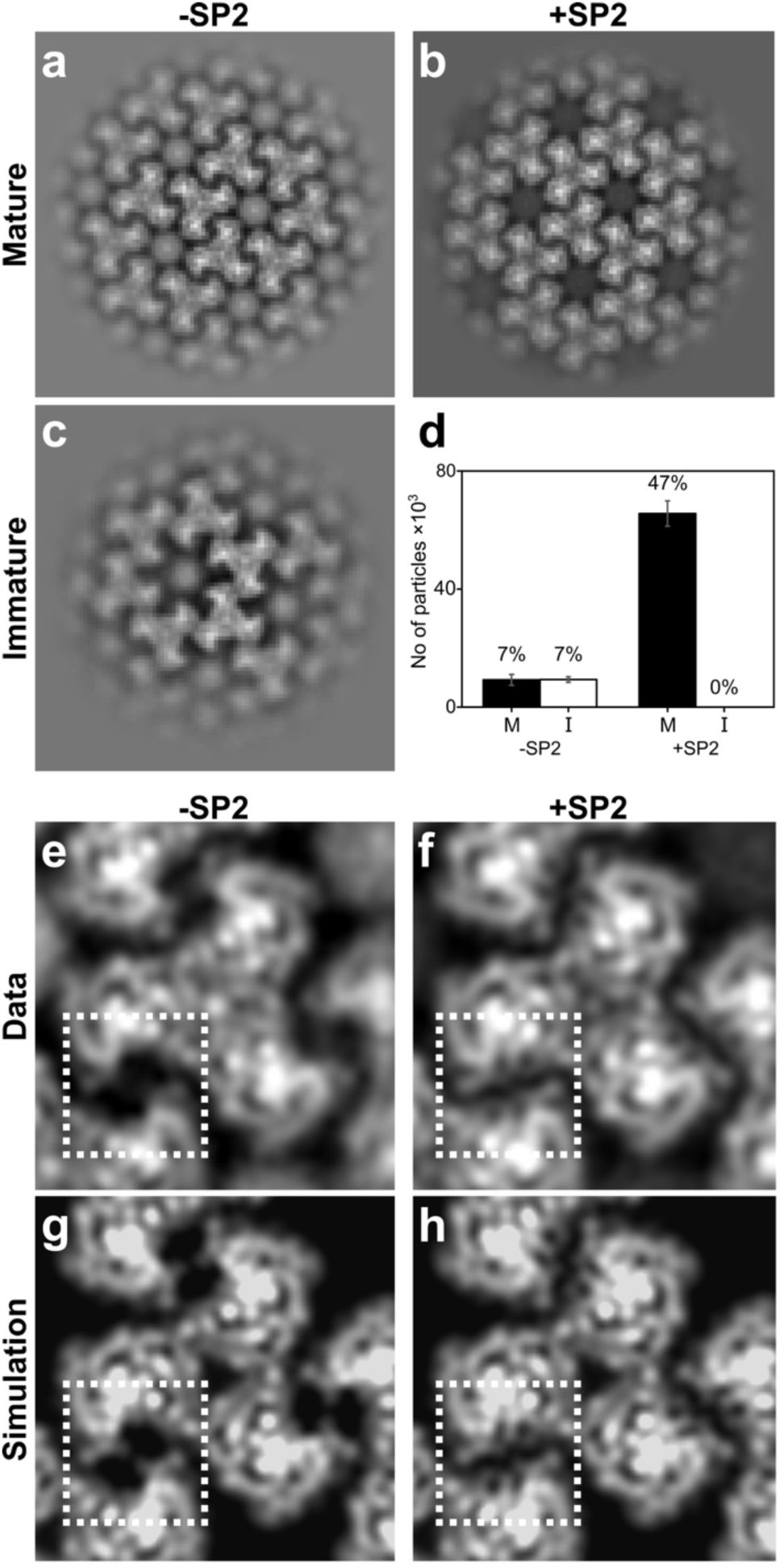
2D Crystallography of HIV-1 MA shows SP2 binding in vitro. **a-c,** Example 2D class averages of MA assembled on lipid monolayers with or without addition of SP2 showing mature (a, b) and immature (c) arrangements. **d,** Quantification of particles that were identified as either mature “M” or immature “I” matrix lattices by 2D classification. 139,500 particles were picked initially for each experiment. Error bars indicate standard deviation between three experiments. **e, f,** An extra density in the side binding pocket of MA highlighted by the dashed square is visible in the +SP2 2D class average (f). **g, h,** Simulated projections of cryo-EM density map of the matrix from MASP1 HIV-1 cleavage mutant. The side pocket density corresponding to SP2 was removed in the structure used to generate (g) to simulate the absence of the SP2 peptide

We generated high-resolution 2D class averages for the mature-like MA lattices formed in the presence or absence of SP2 (Fig. 4e,f), and compared these to simulated projections of MA structures containing or lacking SP2 (Fig. 4g,h). Mature-like MA lattices formed in the presence of SP2 contain an additional density in a position exactly corresponding to that of SP2 in the cryo-EM structure. Together, these results demonstrate that SP2 can induce formation of mature, membrane-bound MA lattices by binding to the described cleft in the absence of any other components.

## Discussion

It has been an enduring question what role the highly conserved SP2 peptide and the two flanking cleavage sites play in the virus life cycle. Our data now indicate that after its proteolytic release from Gag, SP2 serves as the trigger for MA lattice maturation. Our high-resolution, in-virion structure of the MA lattice reveals that after binding a pocket in MA, SP2 becomes an integral component of the inter-trimeric contacts that form the mature MA lattice. These data contrast with the previous model, which proposed that MA maturation is coupled to binding of the same pocket by the lipid PI(4,5)P_2_, which would thereby become partially extracted from the lipid bilayer ^22^.

In the previous model for MA maturation, PI(4,5)P_2_ extraction provided an explanation for reported changes in the mechanical properties of the virus upon maturation ^24,25^. The data presented here place the N-terminal residues of SP2 directly at the viral envelope in a position likely to directly interact with the inner leaflet. The N-terminal residues include lysines, which could interact with lipid headgroups, and hydrophobic residues, which could insert into the membrane bilayer. This positioning would allow SP2 to modulate the properties of the viral membrane and thereby explain the reported changes in virus mechanical properties upon MA maturation.

The highly conserved SP2 sequence (Fig. 3c), the two strictly conserved SP2-flanking PR cleavage sites, and the structure and stability of the mature MA lattice, indicate that SP2 cleavage and MA maturation carry a distinct replicative advantage for the virus. The sequence conservation of the SP2-encoding region can be partly accounted for by coding requirements – this part of the HIV-1 genome includes two overlapping reading frames (*gag* and *pol*) and must maintain an RNA hairpin that promotes ribosomal frameshifting ^33^. There is also some evidence that, prior to proteolytic cleavage, interactions between SP2 and NC may contribute to fine-tuning of NC-nucleic-acid binding ^34-36^. We find that, in addition, release of SP2 and maturation of MA correlate with the virus gaining full fusion competence. However, the HIV-1 mutant NC-SP2, which is defective in release of SP2, does not display an obvious infectivity defect in standard cell culture assays ^31,34,37^. We hypothesize that slower fusion kinetics when MA is in an immature state may be limiting in primary cells with lower receptor and co-receptor levels, thus providing an evolutionary advantage to fast fusion. The generation of a highly structured and conserved mature matrix layer suggests that mature MA could also play other roles in the post-maturation stage of the HIV-1 replication cycle, potentially regulating early post-entry events. The full functional relevance of MA maturation for efficient HIV-1 spread and pathogenesis will require studies in complex systems more closely related to *in vivo* infection conditions.

The pocket where SP2 binds to MA is versatile in its specificity, being able to bind tRNA as well as PI(4,5)P_2_ and SP2 ^11,23,38^. Considered together with our observations, this suggests that exchange of ligands in this pocket determines the changing functions of MA during the viral replication cycle. In the cytosol of the virus-producing cell, the pocket is bound by tRNAs. tRNAs are displaced from the pocket upon binding of MA to the lipid bilayer during virus assembly. Subsequent binding of the pocket by cleaved SP2 then induces MA rearrangement upon maturation in the released virion.

SP2-mediated MA maturation shows intriguing parallels with the regulation of CA lattice maturation by the other Gag spacer peptide, SP1. SP1 is a structural component of the immature Gag lattice, and during maturation, release of SP1 by proteolytic cleavage promotes transition to the mature CA lattice ^8,39,40^. HIV-1 thus has evolved two spacer peptides in its structural polyprotein Gag, which through proteolytic cleavage at the final steps of Gag processing, serve central functions in the conversion of the immature virus particle into the mature, infection-competent virion. SP1 stabilizes the immature Gag lattice and its proteolytic cleavage promotes formation of the mature, cone-shaped capsid; proteolytic cleavage of SP2 promotes formation and stabilization of the mature matrix lattice (Extended Data Fig. 14).

## Materials and Methods

### Plasmids

All plasmids were based on the subviral plasmid pcHIV ^41^ which encodes all proteins of HIV-1_NL4-3_ except for Nef. The pcHIV variants MA-SP1, NC-P6, MA-SP1-NCP6, MA-NC, PR-have been described before ^7,37,42^. MA-SP2 was created by introducing mutations at the NC-SP2 cleavage site ^31^ into pcHIV(MA-NC) by overlap PCR using oligos 5’GAGAGACAGGCTTCTTTTTTAGGGAAGACCTGGCCTTCCCACAAGGG 3^‘^ and 5^‘^CCCTTGTGGGAAGGCCAGGTCTTCCCTAAAAAAGAAGCCTGTCTCTC 3^‘^.

### Cell lines and virus particle production

HEK293T cells were grown Dulbecco’s modified Eagle’s medium, 100 U/ml penicillin, 100 μg/ml streptomycin, and 10% fetal calf serum (FCS). At 80% confluency, cells were split 1:3 into T175 flasks (CELLSTAR®, Greiner BIO-ONE) the day before transfection. Cells were transfected with pcHIV (70µg/T175 flask) in three T175 flasks per variant using a standard calcium phosphate transfection procedure. At 48 h post transfection, tissue culture supernatant was collected and cleared through a 0.45μm pore filter. The filtered supernatant was layered on top of a 20%(w/v) sucrose cushion and subjected to ultracentrifugation at 107000g for 1.5h at 4C. The pellet was resuspended in phosphate buffered saline (PBS) and stored in aliquots at -80 °C. For quantification, particle associated reverse transcriptase activity was determined using the Sybr Green Product Enhanced Reverse Transcription assay (SG-PERT) ^43^.

### Immunoblotting

Particles were separated by SDS-PAGE (20%; acrylamide: bisacrylamide 30:1). Proteins were transferred to a nitrocellulose membrane (Millipore) by semidry blotting and stained with the indicated antisera in PBS/ Intercept^®^ (PBS) Blocking Buffer (LICORBIO^TM^) (sheep αCA, polyclonal, 1:5,000 [in-house]; rabbit αNC, polyclonal, 1:400 [in-house], followed by corresponding secondary antibodies IRDye in PBS/ Intercept^®^ (PBS) Blocking Buffer (LICORBIO^TM^) (Donkey anti-sheep, 1:10,000 [LiCOR Biosciences]; and Donkey anti-rabbit, 1:10,000 [Rockland]). Detection was performed using a Li-COR Odyssey CLx infrared scanner (Li-COR Biosciences) according to manufacturer’s instructions. Blots are shown in Extended Data Fig. 2a.

### Cryo-EM

All cryo-EM samples of purified HIV-1 particles were prepared and imaged similarly. Purified virus was diluted in PBS buffer in 1:3 (v/v) ratio. 3 ml of the diluted virus sample was applied on a glow-discharged Quantifoil 2/2 holey carbon grid, Cu 300 mesh (Quantifoil Micro Tools GmbH) and plunge-frozen into an ethane-propane 1:1 mixture using the Leica EM GP2 (100% humidity, blot time 3.5 s, 20 °C). Grids were loaded into Titan Krios G4 transmission electron microscope operated at 300 kV, equipped with CFEG electron source, Falcon4i direct detector camera and Selectris X energy filter (ThermoFisher Scientific). Images were collected in electron-event representation (EER) format ^44^ using EPU (v3.0) (ThermoFisher Scientific). All datasets were collected at a magnification of 135,000x, resulting in a pixel size of 0.95 Å, with acquisition times ranging from 3.75 to 4.15 seconds and with a total dose of 40 e^-^/Å^2^. Detailed data acquisition parameters and the number of micrographs for all datasets are given in Extended Data Table 1.

EER movies were rendered as an 8k x 8k grid and further Fourier cropped into a 4k x 4k grid using RELION-4.0 ^45^. The movies were motion-corrected, dose-weighted, and averaged using RELION-4.0 MotionCorr2 algorithm ^46^. Frames were dose-fractionated into groups resulting in a dose of 0.8 e^-^/Å^2^ per fraction. CTF estimation was performed using the patchCTF algorithm in cryoSPARC 4.0 ^47^. Particle picking was performed using crYOLO (v1.9.2) ^48^. For each dataset, a new model was trained in crYOLO using a training dataset annotated in a randomly selected set of 50-100 micrographs. Annotation was performed in the crYOLO boxmanager GUI placing positions all over the visible surface of an HIV virus particle. The picks did not distinguish individual proteins or membranes.

### Cryo-EM data processing PR-mutant

The data processing pipeline for PR-MA is summarised in Extended Data Fig. 3. 9,942 motion-corrected micrographs and 3,568,755 particle positions were imported into cryoSPARC v4.0 ^47^. Particles were extracted with a box size of 480 x 480 pixels, Fourier cropped to 180 x 180 pixels and subjected to several rounds of 2D classification. The first step was to perform a reconstruction of the CA layer. Classes showing side and top views of the capsid layer were selected (1,630,811 particles) and subjected to a heterogeneous refinement with 8 classes, C6 symmetry imposed, and using a previously solved immature structure of an *in-vitro* assembled HIV capsid (EMD-3782) as a starting reference ^49^. Classes showing resolved secondary structures in the capsid layer with visible densities representing matrix and membrane layers were selected (805,005 particles) and reextracted with a box size of 480 × 480 pixels, and Fourier cropped to 416 × 416 pixels. Duplicate particles were removed based on a spatial separation and the accepted particles (676,036 particles) were subjected to three rounds of 3D refinement with local spatial and angular searches (local refinement), C6 symmetry imposed, and a mask comprising the capsid. Local and global CTF refinements were performed in between the 3D refinements. Afterwards, the particles were imported to Relion 4.0, reextracted with a box size of 480 × 480 and Fourier cropped to 416 × 416 pixels. The particles were then subjected to Bayesian polishing in Relion 4.0 ^50^. The polished particles were imported back to cryoSPARC and subjected to two rounds of local refinement.

The coordinates of the CA layer were then used to predict initial coordinates for the MA layer. To do this, the particles from the CA reconstruction were symmetry expanded using 6-fold symmetry and the 3D coordinates of the centre of the box were shifted to the centre of the 3-fold axis of the matrix layer. The particles (3,920,498 particles) were then extracted using the new box centre with a box size of 512 × 512 px and Fourier cropped to 256 × 256 px. Duplicate particles were removed and the accepted particles were reconstructed with C3 symmetry. The resulting map was then used to subtract densities corresponding to the capsid layer from the particles. The subtracted particles were subjected to 3 rounds of heterogeneous refinement in cryoSPARC, with C3 symmetry imposed and 8 classes using a cryo-ET derived reconstruction of the immature matrix lattice (EMD-13087) low pass filtered to 10 Å as a starting reference ^22^. Particles belonging to classes which showed aligned matrix and membrane layers were selected in each iteration. Selected particles (120,104 particles) were then subjected to non-uniform refinement ^51^ with C3 symmetry and a mask comprising both the matrix and membrane layers followed by a local 3D refinement with a mask comprising only the matrix layer.

Next, the dataset was expanded by using the calculated positions of MA trimers to predict the positions of neighbouring MA trimers (lattice expansion). To do this, the particles were symmetry expanded with C3 symmetry, the centre of the box was shifted to a neighbouring matrix trimer, and reextracted with a box size of 512 × 512 px and Fourier cropped to 256 × 256 px. Duplicate particles were then removed. Two iterations of the lattice expansion were then performed. The accepted particles (819,672 particles) were subjected to two rounds of 3D classification without angular search, using 10 classes, C1 symmetry, a focus mask comprising the MA layer and simple initialization mode (the initial volumes were generated from randomly selected particle subsets). Particles were lattice expanded as described above between the two 3D classifications. Classes showing a well resolved MA layer were selected (174,245 particles) and subjected to a final round of local refinement, imposing C3 symmetry and mask comprising the MA layer.

### Cryo-EM data processing MA-SP1

The data processing pipeline for MA-SP1 MA is summarised in Extended Data Fig. 4. 14,222 motion-corrected micrographs and 5,704,512 initial particle positions were imported into cryoSPARC v4.0 ^47^. Particles were extracted with a box size of 512 x 512 pixels, Fourier cropped to 192 x 192 pixels and were then subjected to two rounds of 2D classification. The first step was to perform a high-resolution reconstruction of the CA layer. Classes showing side and top views of the capsid layer were selected (2,969,039 particles) (Extended Data Fig. 4b). To speed up computation, selected particles were split into four roughly equally sized subsets (∼750,000 particles), each of which were subjected to heterogenous refinement with 6 classes, enforced C6 symmetry, using a previously determined structure of an *in-vitro* assembled HIV-1 capsid (EMD-3782) as a starting reference (Extended Data Fig. 4c) ^49^. The highest quality classes from each batch were then pooled and subjected to non-uniform 3D refinement before particle reextraction with a box size of 512 x 512 pixels, Fourier cropped to 384 x 384 pixels. Duplicates particles were removed based on a spatial separation distance and the accepted particles (885,809 particles) subjected to 3D refinement with local angular and spatial searches (local refinement), imposed C6 symmetry, and a mask comprising the CA layer. The particles were then subjected to local CTF refinement, followed again by 3D refinement. Particles were re-extracted with a box size of 512 x 512 pixels, and Fourier cropped to 450 x 450 pixels, and again subjected to Local CTF refinement and local 3D refinement. Heterogenous refinement was then performed to remove any remaining low-quality particles, using 3 classes, from which the highest quality class was selected. Global CTF refinement ^52^, and further local refinement was then performed. Afterwards, the particles were imported to Relion 4.0 and subjected to Bayesian polishing ^50^. The polished particles were imported back to cryoSPARC and subjected to further 3D refinement and local CTF refinement to generate a final high resolution immature CA reconstruction (Extended Data Fig. 4d).

The coordinates of the CA layer were then used to define initial coordinates and orientations for the MA layer focused reconstruction. The 3D coordinates of the centre of the box were shifted to the centre of the 3-fold symmetry axis of the matrix layer, which could be seen in the high-resolution CA reconstruction. Local refinement, without provision of a new reference, was then performed with imposed C3 symmetry and a mask comprising only the MA layer (Extended Data Fig. 4e). The refined particles were then lattice expanded, with the centre of the box shifted to the neighbouring 6-fold symmetry axis, before duplicate particles were removed based on a spatial separation distance. The accepted particles (1,398,844 particles) were subjected to a further round of local refinement with C6 symmetry enforced. The particles were once again lattice expanded, with the centre of the box shifted to the neighbouring matrix timers. Duplicate particles were removed and the accepted particles (5,486,693 particles) were reconstructed with C3 symmetry imposed to generated the final MA reconstruction (Extended Data Fig. 4e).

### Cryo-EM data processing MA-NC

The processing strategy for MA-NC was almost identical to that of MA-SP1, for CA and the subsequent MA refinement steps. The data processing pipeline is summarised in Extended Data Fig. 5.

### Cryo-EM data processing WT

The data processing pipeline for WT MA is summarised in Extended Data Fig. 6. 19,530 motion-corrected micrographs and 5,801,053 initial particle positions were imported into cryoSPARC v4.0 ^47^. Particles were initially extracted with a box size of 480 x 480 pixels, Fourier cropped to 240 x 240 pixels and subjected to two rounds of 2D classification. Classes showing top-views and side-views of the MA layer were selected (Extended Data Fig. 6b) and subjected to two rounds of heterogenous refinement, each with 4 classes and imposed C6 symmetry, using a cryo-ET derived mature MA lattice reconstruction as a starting reference (EMD: 13088) (Extended Data Fig. 6c). The resulting highest quality class was selected (61,672 particles) and subjected to non-uniform 3D-refinement with C6 symmetry imposed^51^. Duplicate particles were removed based on spatial separation distance and the accepted particles (57,260 particles) were then subjected to 3D-refinement with local spatial and angular searches (local refinement), using a refinement mask comprised of the MA layer. The particles were lattice expanded, with the centre of the box shifted to the neighbouring 6-fold symmetry axis, and re-extracted with a box size of 480x480 pixels, Fourier cropped to 356 x 356 pixels (276,547 particles). The particles were subjected to local CTF refinement, followed by a further round of local refinement. The particles were once again lattice expanded, with the centre of the box was shifted to the neighbouring matrix timers. Duplicates were again removed and the accepted particles (898,502 particles) were finally reconstructed with imposed C3 symmetry (Extended Data Fig. 6d).

### Automated lipid fitting

Automated docking of lipid candidates into the MA-SP1 ligand density (Cholsterol, PI(4,5)P_2_, Phosphatidyl Serine (PS), Phosphatidyl Choline (PC) and Phosphatidyl Ethanolamine) was performed using RosettaEmerald, using the protocol described in Muenks et al 2023 ^53^. All resulting fits were visually inspected and Q-scores of all ligand fits were determined using MapQ in USCF chimera ^54^. The top ten fits for each ligand, according to Q-score, are provided (Extended Data Fig. 7).

### Fourier analysis of 2D class averages of cleavage mutants

Cryo-EM data preprocessing and particle picking were performed as described in the cryo-EM subsection. Images were extracted with a box-size of 480 × 480 px and downsampled to 240 × 240 px resulting in a final pixel size of 1.9 Å. 2D class averages of mutants that displayed side-views of membranes regardless of the presence of matrix or capsid layer were selected manually in cryoSPARC v4.11. Afterwards, the 2D classes were reoriented according to a reference class where the membrane bilayer was oriented perpendicularly to the y-axis of the 2D class using cross-correlation (Extended Data Fig. 8). Then, the pixel values along the matrix layer (section parallel to the inner leaflet of the viral membrane) were interpolated with 1024 points and exported as one-dimensional vectors. The vector was filtered using a Hann function and zero-padded to a total of 4096 sampling points. Fast Fourier transformation (FFT) of the zero-padded signal was plotted in MATLAB v2022a (MathWorks) and analysed for peaks corresponding to matrix lattice spacing frequencies. All the signal processing steps were performed in MATLAB v2022a (MathWorks).

### Model building and refinement of MA-SP1 MA-SP2 atomic model

The solution structure of myristoylated HIV-1 MA (PDB:2H3I) ^11^ was used an initial MA structure for building into the MA-SP1 density. Initial co-ordinates for SP2_11-16_ were generated using AlphaFold (v2.2.0) ^55^, which were fit roughly into the SP2 density in USCF Chimera ^54^. The initial model was then flexibly fit, with manual adjustments, into the density with ISOLDE (ChimeraX) ^56,57^. A single round of real space refinement was then performed in Phenix-1.21. Final model validation statistics and the map-to-model FSC were calculated in Phenix-1.21 ^58^ and are given in Extended Data Table 2.

### ModelAngelo Predictions of SP2

The MASP1 cleavage mutant mature matrix map with a resolution of 3.1 Å was used for automated ModelAngelo ^59^ atomic model predictions. The box size was cropped to 128 x 128 px and only sequences built into the central trimer were considered. The ModelAngelo job was run with default parameters in Relion-5.0. The sequence input was either the HIV-1 Gag sequence or no sequence. Afterwards, sequences built into the side pocket cryo-EM densities from the central MA trimer were extracted and analysed by Clustal Omega multiple sequence analysis tool ^60^. The ModelAngelo automated model building and sequence prediction results are shown in (Extended Data Fig. 10).

### Expression and purification of HIV-1 matrix

The expression plasmid pET11b-MA encodes HIV-1 pNL4–3 matrix domain with 6 C-terminal His tag ^61^. BL21(DE3) *E. coli* competent cells for protein expression were co-transformed with the pET11b-MA plasmid and plasmid encoding for the yeast N-terminal myristoyltransferase. The protein was expressed and purified as described previously ^16,62^ with modifications. Cells were grown at 37°C at 180 rpm in a lysogeny broth (LB) medium containing 100 mg/l of Ampicylin and 50 mg/l of Kanamycin. When A_600_ reached ∼0.6; 15 mg/l of myristic acid (Sigma-Aldrich) was added to the LB medium. After 30 minutes, cells were induced with 0.5 mM isopropyl β-D-1-thiogalactopyranoside and grown for another 5 hours at 37°C at 180 rpm. Afterwards, the cells were spun down at 3500×g and the pellets were stored at -80°C until further use. For purification, 6 grams of the cell pellet was diluted in 60 ml of lysis buffer (25 mM Tris pH 8, 500 mM NaCl, 2 mM TCEP, and 2 mM PMSF) and sonicated for 2 minutes. Then, 20 µl of benzonase was added to the lysed cells. The lysate was incubated on ice for 10 minutes then spun down at 50,000 rpm at 10°C for 45 minutes (Beckman Coulter, 50.2 TI rotor). The supernatant was collected and treated with polyethyleneimine to a final concentration of 0.03% incubated on ice for 5 minutes and subsequently centrifuged at 10,000 rpm at 4°C for 10 minutes (Beckman coulter, JA-25.50 rotor). The supernatant was collected and powdered ammonium sulfate (approx 15 g) was added to the supernatant on ice with constant stirring until protein precipitate was observed followed by centrifugation at 10,000 rpm at 4°C for 10 minutes (Beckman coulter, JA-25.50 rotor). The pellet was resuspended in 4 ml of binding buffer (20 mM Tris-HCl, pH 8, 100 mM NaCl, 2 mM TCEP) and loaded onto a HisTrap column (Cytiva) equilibrated with the binding buffer. The column was then washed with a wash buffer (20 mM Tris-HCl, pH 8, 100 mM NaCl, 2 mM TCEP, 20 mM imidazole) and the protein was subsequently eluted with elution buffer (20 mM Tris-HCl, pH 8, 100 mM NaCl, and 2 mM TCEP, 250 mM imidazole). Fractions containing myrMA were collected and further purified by gel filtration using Superdex 75 16/600 (Cytiva) equilibrated in buffer containing (20 mM Tris-HCl, pH 8, 500 mM NaCl, 1 mM TCEP). Fractions containing myrMA were collected and stored at -80°C. The presence of myristoylation modification was confirmed by mass spectrometry.

### 2D crystallisation of HIV matrix

2D crystallisations were performed in a cleaned polytetrafluoroethylene (PTFE) block containing 60 ml side-entry reservoirs, combining previous protocols ^63^). 58 ml of crystallisation buffer (12 mM sodium phosphate buffer pH = 7.8; 2.5 mM sodium acetate pH = 7.6; 150 mM sodium chloride; 10% glycerol) ^32,64^ was added to 6 crystallisation reservoirs. Afterwards, 1 µl of a freshly prepared lipid mixture mimicking the inner leaflet of the viral membrane (molar fractions: 31% cholesterol; 6% POPC; 29% POPE; 27% POPS; 7% PI(4,5)P_2_ ^65^) in 9:1 (v/v) chloroform:methanol solution at a lipid concentration of 0.01 mg/ml was carefully added on top of each buffer surface. The PTFE block was then incubated in a closed petri dish with a wet filter paper placed underneath the block for 60 minutes to allow a lipid monolayer to form at the air-water interface. Then a Quantifoil 2/2 holey carbon grid, Au 200 mesh (Quantifoil Micro Tools GmbH) was placed on top of each reservoir. A solution containing purified myristoylated HIV-1 matrix protein (myrMA) was then injected into each reservoir from the side entrance. The final concentration of myrMA in the reservoir was 12 mM. After 10 minutes, HIV-1 spacer peptide 2 (SP2) in PBS buffer was added to 3 of the 6 experimental reservoirs to a final concentration of 120 mM. The same amount of PBS buffer was added to the 3 control reservoirs. All samples were incubated for an additional 60 minutes, grids were carefully lifted from the surface of the reservoirs, and plunge-frozen using Vitrobot Mark IV (4s blot time; 3 blot force; 100% humidity).

Grids were loaded into a Talos Glacios cryo-transmission electron microscope operating at 200 kV and equipped with a Falcon4i direct electron detector (ThermoFisher Scientific). For each grid, 5 grid squares were selected manually and in each square 30 holes were randomly selected in the EPU software (v3.0). A single acquisition position was selected in the centre of each hole resulting in 150 micrographs automatically collected from each grid. The micrographs were collected as movies of 40 frames with a total dose of 40 e^-^/ Å^2^ at a magnification of 92,000x resulting in a pixel size of 1.20 Å per pixel. Movies were motion-corrected, dose-weighted, and averaged using the RELION-4.0 MotionCorr2 algorithm ^45,46^. CTF estimation was performed using CTFfind4 ^66^. Particles were picked as a grid of points separated by 128 pixels placed in a 4k x 4k micrograph resulting in 139,500 particles for each dataset (Extended Data Fig. 13b). Particles were extracted with a box size of 256 px and Fourier cropped to 128 px. The particles were then imported to cryoSPARC v4.0 ^47^ and subjected to two rounds of 2D classification (Extended Data Fig. 13). Classes showing 2D crystal matrix lattice were selected and particles from these classes were considered as particles containing a 2D crystal lattice. All 6 datasets were processed the same way.

### Plasmids, reagents, and cell lines for fusion assays

HIV-1 GagPol was expressed by pCMV ΔR8.2 (Addgene plasmid #12263). The pCAGGS HIV-1_JRFL_ gp160 expression plasmid was kindly provided by Dr. James Binley. The pN1 CypA-HiBiT plasmid (made by Dr. Jonathan Grover) was derived from pEGFP-N1-CypA. To generate this plasmid, human CyclophilinA was cloned from HeLa cDNA and inserted into pEGFP-N1 using the EcoRI and BamHI sites. To generate CypA-HiBiT, a synthetic oligo, which encoded the linker sequence “GSGSSGGGGSGGGGSSG” followed by the HiBiT peptide “VSGWRLFKKIS,” was inserted to replace EGFP at the C-terminus of CypA using the BamHI and NotI sites. This construct is based on a similar design by Dr. Gregory Melikyan.

The Gag cleavage mutants were constructed via site directed mutagenesis and Gibson assembly. The mutations for pCMV ΔR8.2 MA-CA, pCMV ΔR8.2 MA-SP1, pCMV ΔR8.2 MA-NC, and pCMV ΔR8.2 MA-p6 were recreated from previous literature ^5,67,68^. Gene blocks of the MA-CA, MA-SP1, MA-NC, and MA-p6 GagPol sequences were ordered from Twist Bioscience (San Francisco, CA) and combined with fragments of the pCMV ΔR8.2 backbone amplified by PCR for Gibson assembly using the Gibson Assembly® Master Mix from New England Biolabs (Ipswich, MA). The mutations for pCMV ΔR8.2 MA-SP2 and pCMV ΔR8.2 NC-p6 were created using site directed mutagenesis using the pCMV ΔR8.2 MA-p6 and pCMV ΔR8.2 constructs as templates, respectively. These fragments were combined with pCMV ΔR8.2 backbone fragments amplified by PCR and combined with Gibson assembly using the same protocol as above.

The human CD4-expressing vector pcDNA-hCD4 was kindly provided by Dr. Heinrich Gottlinger. The pMX-puro PH-PLCΔLgBiT plasmid was kindly provided by Dr. Zene Matsuda ^69^. The following reagents were obtained through the NIH HIV Reagent Program, Division of AIDS, NIAID, NIH: Indinavir sulfate, ARP-8145 and Ritonavir, ARP-4622, both contributed by DAIDS/NIAID; Human CCR5 expression vector (pcCCR5), ARP-3325 contributed by Dr. Nathaniel Landau.

HEK293E cells were grown in the presence of 5% CO_2_ using RPMI-1640 media from Thermo Fisher Scientific (Waltham, MA) supplemented with 10% fetal bovine serum (FBS), 100 U/mL of a penicillin and streptomycin solution, and 2 mM L-glutamine. Cells were transfected at 60%–80% confluency and culture medium was exchanged before transfection.

### Virus-like particle preparation for fusion kinetics

Virus-like particles (VLPs) were produced by transfecting three 10 cm plates of HEK293E cells with 12 µg DNA per 10 cm plate using polycation polyethylenimine (PEI) (pH 7.0, 1 mg/mL). Immature VLP preparations were transfected with a final concentration of 1 µM Indinavir and 10 µM Ritonavir in the cell culture media. Plasmids were transfected in a 1:1:1 ratio of Gag:Env:HiBiT. Cell culture media was collected two days after transfection, then spun down for 5 min to pellet cells. Supernatants were transferred to 38.5 mL ultracentrifuge tubes and underlayed with 5 mL sterile-filtered 15% sucrose in PBS. VLPs were then pelleted by ultracentrifugation at a maximum of 131,453 rcf using a Beckman Coulter (Brea, CA) SW28 swinging bucket rotor at 27,000 rpm for 1 hour at 4°C. Supernatant was removed and viruses were resuspended in 1:100 volume (300 µL) of neat CO_2_-independent media (Thermo Fisher Scientific; Waltham, MA). Particles produced in the presence of protease inhibitors were resuspended in neat CO_2_-independent media with final concentrations of 1 µM Indinavir and 10 µM Ritonavir. Each VLP prep was aliquoted into ∼20 tubes at 15 µL per tube and stored at -80°C until use. After harvesting, each aliquot of VLPs was used once for fusion kinetics assays to minimize particle destruction during freeze-thaw cycles.

### VLP normalization for fusion kinetics

VLP volumes were normalized on HiBiT incorporation using the Nano-Glo® HiBiT Lytic Detection System from Promega (Madison, WI) according to the manufacturer’s instructions. Three volumes of the VLPs (2 µL, 4 µL, and 6 µL) were measured per sample. Plates were placed in a Promega GloMax Explorer GM3500 Multimode Microplate Reader and read using the manufacturer’s suggested protocol. Readings were then graphed in Excel with a linear trendline. The trendline equation was used to calculate the volume of each sample containing 3×10^7^ RLU of HiBiT.

### Fusion kinetics assay

The split nanoluciferase fusion kinetics assay was based on the assay described in Yamamoto *et al.* ^69^ Briefly, HEK293E cells were transfected with a 1:1:1 ratio of CD4, CCR5, and PH-LgBiT. After 24 hours, cells were harvested and Endurazine (Promega; Madison, WI) and DrkBiT peptide were added to the solution to a final concentration of 1x and 1 µg/mL, respectively. A white 96-well flat-bottom plate was prepared with 100 µL of the cell solution per well. Prepared VLPs were added to the wells and spinoculation was performed at 1200 rcf and 12°C for 2 hrs. The plate was read using a Promega GloMax Explorer GM3500 Multimode Microplate Reader using an automatic protocol for 24 rounds of reading wells every 2 min with a 1.5 sec exposure time per well. Readings were exported into Excel where RLU was calculated, then RLU was processed into percent of total fusion. The percent of total fusion curves were processed using an in-house Mathematica code (written by Dr. Alexander Lee) to generate T_1/2_ for each sample. Results were plotted using GraphPad Prism.

## Acknowledgements

This work was funded by the Deutsche Forschungsgemeinschaft (DFG, German Research Foundation) Projektnummer 240245660 - SFB 1129 (project 5 HGK, project 6 BM, project 21 JAGB), DFG KR 906/7-1 (HGK), NIH/NIAID R37 AI150560 (WM), and the Max Planck Society (JAGB). E.N. was supported by NIH/NIAID T32 AI055403. Cryo-EM data was collected at MPI Biochemistry. We thank Lisa Regner for plasmid pcHIV(MA-SP1:NC-p6); Hui Guo and Zunlong Ke for assistance with data collection; Zunlong Ke for advice on data analysis; Florian Beck, Jesuraj Rajan Prabu and Inga Wolf for assistance with computing infrastructure; Alexander Lee for assistance with fusion kinetics data analysis.

## Author contributions

J.C.V.S, D.H., E.N., W.M., B.M., H.G.K. and J.A.G.B. designed research; S.D.S and M.A. prepared and analysed virus samples; K.Q. performed preliminary experiments. M.B. prepared protein; R.A.D. provided reagents and advice. J.C.V.S. and D.H. carried out cryo-EM and related data processing; J.C.V.S, D.H. and J.A.G.B analyzed and interpreted structural data; E.N. performed fusion kinetics experiments and analysis; J.C.V.S, D.H., E.N., W.M., and J.A.G.B. designed and prepared figures; J.C.V.S., D.H. and J.A.G.B. wrote the manuscript with input from all authors; B.M., H.G.K., W.M., and J.A.G.B. obtained funding and managed the project.

## Data Availability

Structures determined by electron microscopy are deposited in the Electron Microscopy Data Bank under accession codes EMD-XXXXX. Corresponding molecular models are deposited in the Protein Data Bank under accession codes XXXX. Any additional information required to evaluate the conclusions of the paper is included in the paper or available from the lead author on request.

---

**Extended Data Fig. 1:**
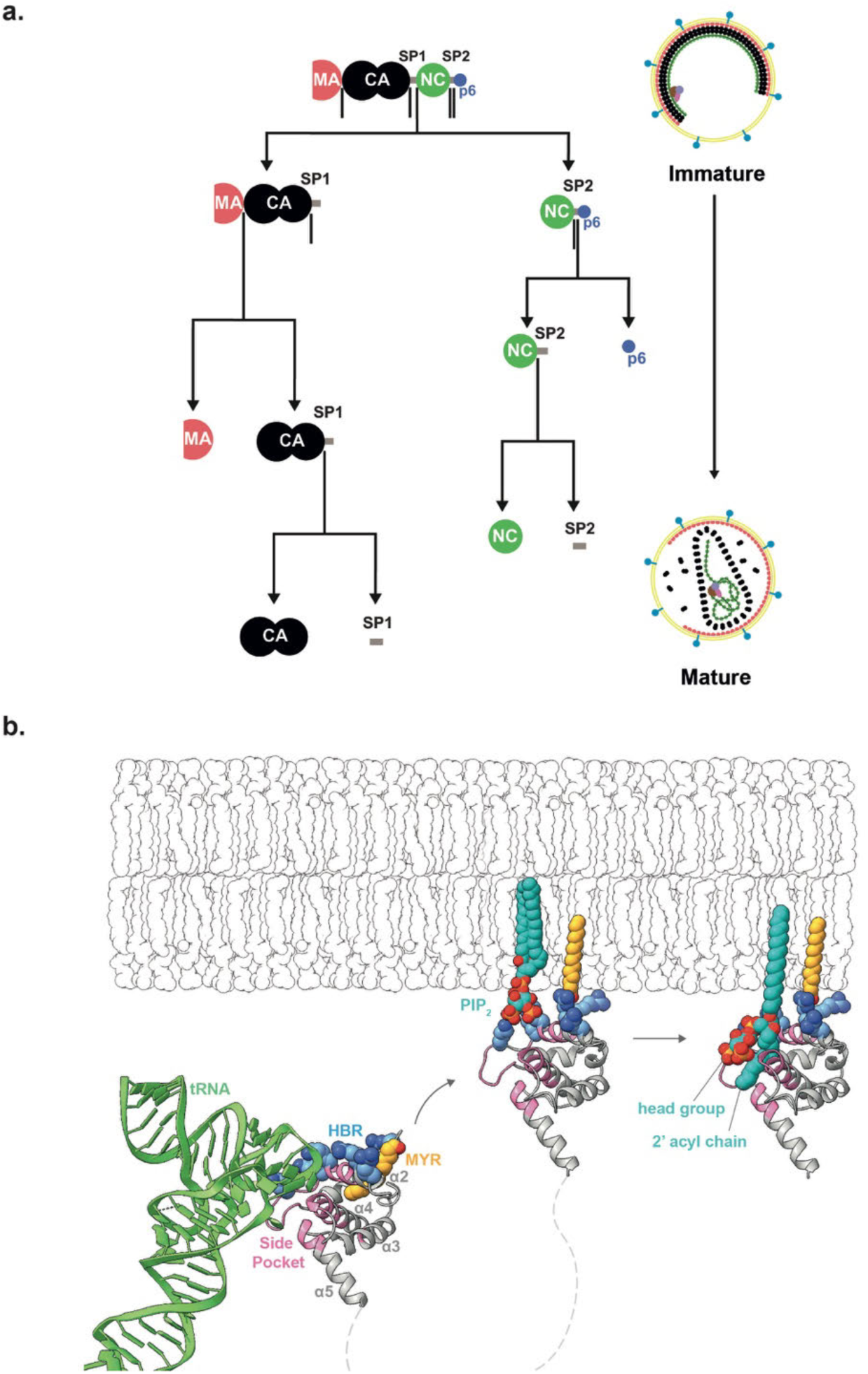
Schematic of the Gag cleavage cascade and current model of MA interactions at the plasma/viral membrane. **a,** Schematic illustration of the Gag proteolytic cleavages that occur during HIV-1 maturation. Cleavages are ordered by relative rate as measured in-vitro ^70^. **b,** MA in the cytoplasm can bind tRNA (green) via the side pocket. MA binds to PIP_2_ (blue lipid) and cholesterol rich domains on the plasma membrane during viral assembly. This binding is facilitated by the insertion of a previously sequestered myristoyl group (yellow) and by electrostatic interactions at the HBR. Bound tRNA sterically prevents membrane binding of MA, thus tRNA^Lys3^ is released upon membrane binding, leaving the side pocket unoccupied. Upon maturation the current model is that that MA partially extracts molecules of PI(4,5)P_2_ from the inner leaflet of the viral membrane, binding the phosphatidylinositol head group and one of the acyl tails within the side pocket. Aspects of this model are directly contradicted by the results of the current study.

**Extended Data Fig. 2:**
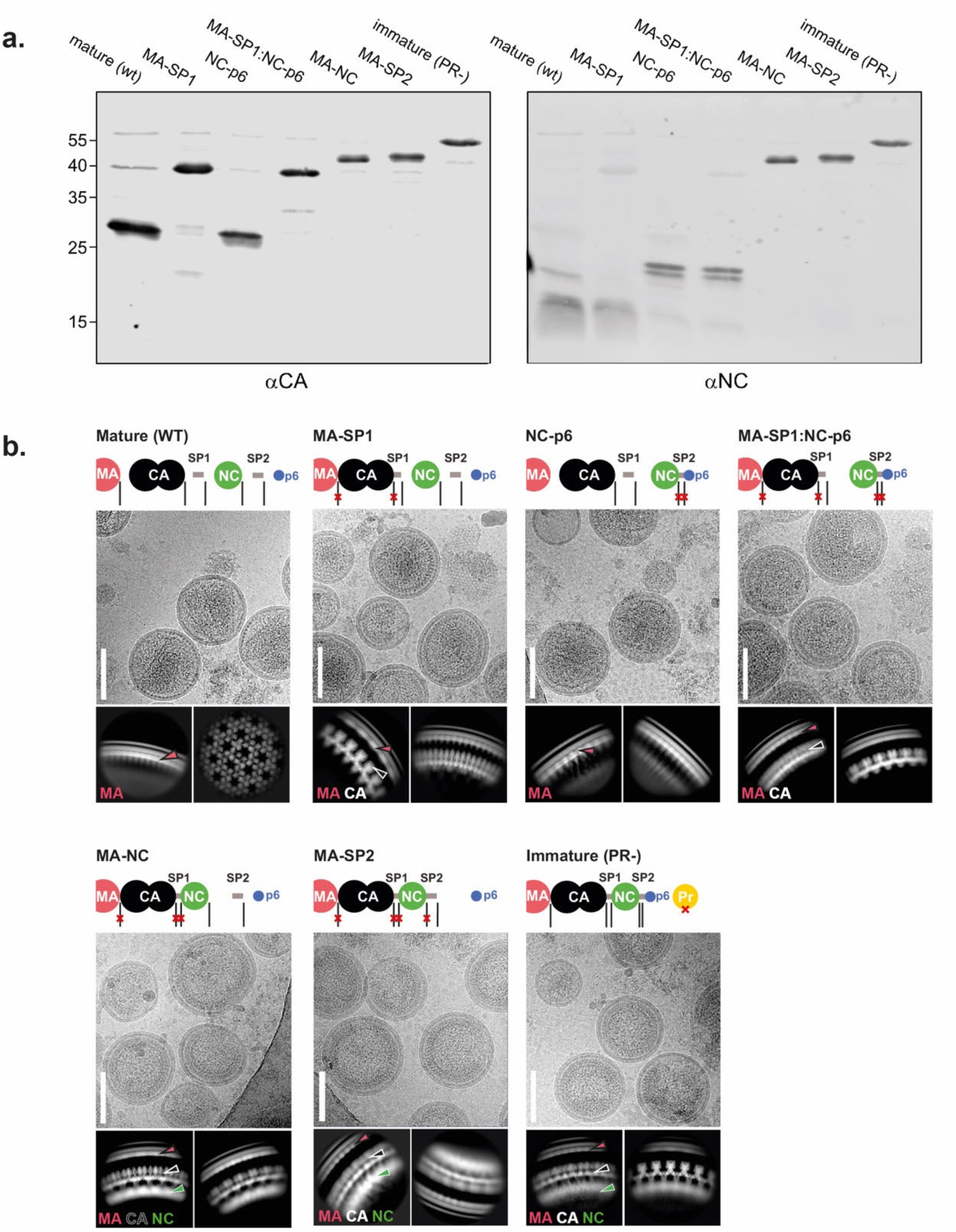
Representative micrographs, 2D classes and western blot analysis of HIV-1 particles used in this study. **a,** HEK293T cells were transfected with pcHIV carrying the indicated Gag cleavage site mutations. Particles were collected at 48 h post transfection by ultracentrifugation. Viral proteins were detected by immunoblotting with the indicated polyclonal antisera. Antibodies were detected using a LI-COR Odyssey Clx infrared scanner, using secondary antibodies and protocols provided by the instrument’s manufacturer. Positions of molecular mass markers are indicated at the left. **b,** For each dataset a representative cryo-EM micrograph is shown, illustrating the overall viral morphology (scale bars: 100 nm). Representative high resolution 2D classes of the Gag layer are also shown, labelled with the features that could be identified, including MA, CA. and NC layers. Schematic illustrations of the processing state of Gag constructs are given for each dataset.

**Extended Data Fig. 3:**
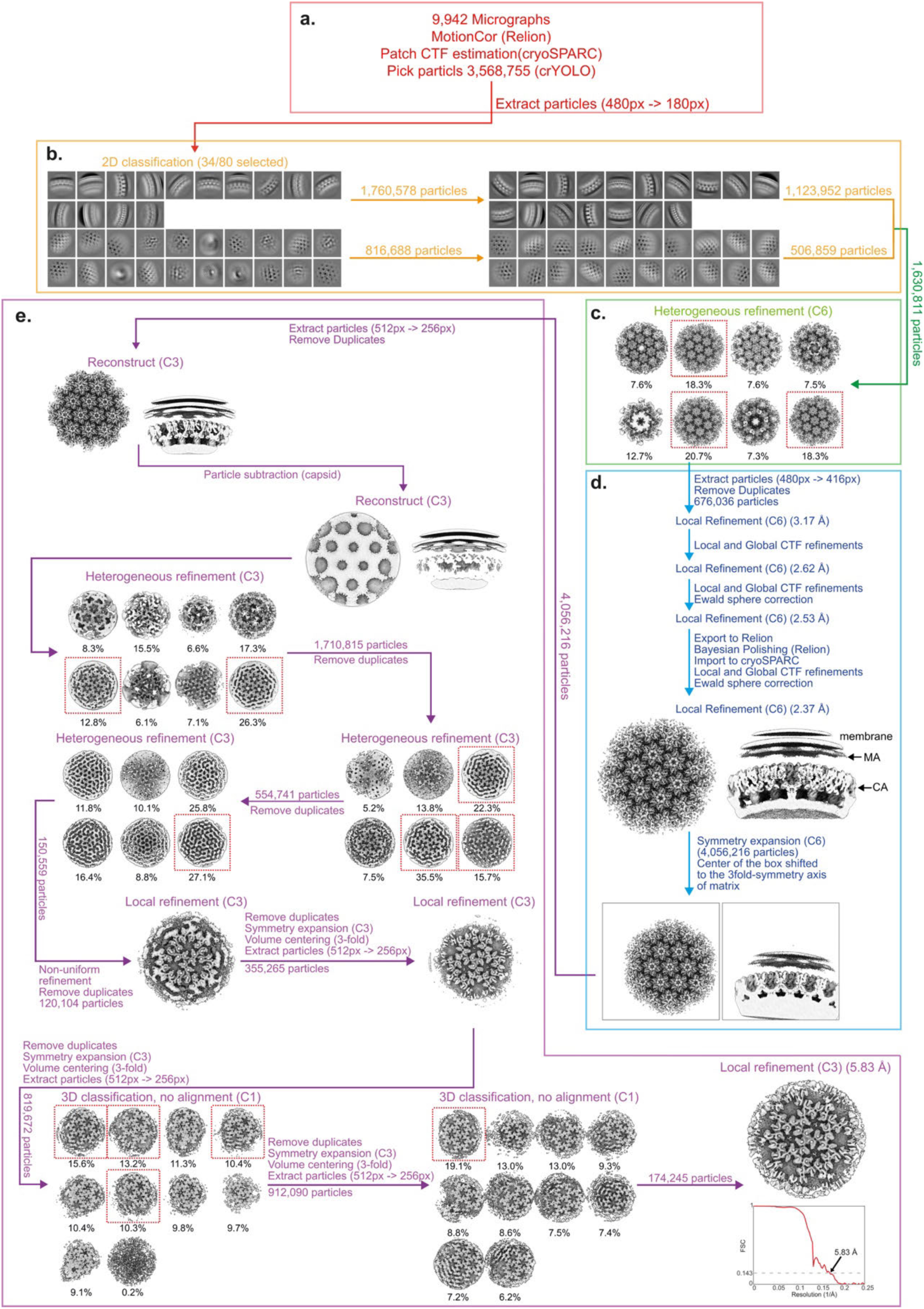
Single-particle cryo-EM processing pipeline for the Immature (PR-) MA lattice. For a detailed description of the steps illustrated see Materials and Methods **a,** EM micrograph preprocessing, particle picking and particle extraction. **b,** 2D classification of the Gag layer. **c,** Heterogeneous refinement of the CA layer. **d,** Heterogeneous refinement and classification of the MA layer. **e,** Local refinement and FSC.

**Extended Data Fig. 4:**
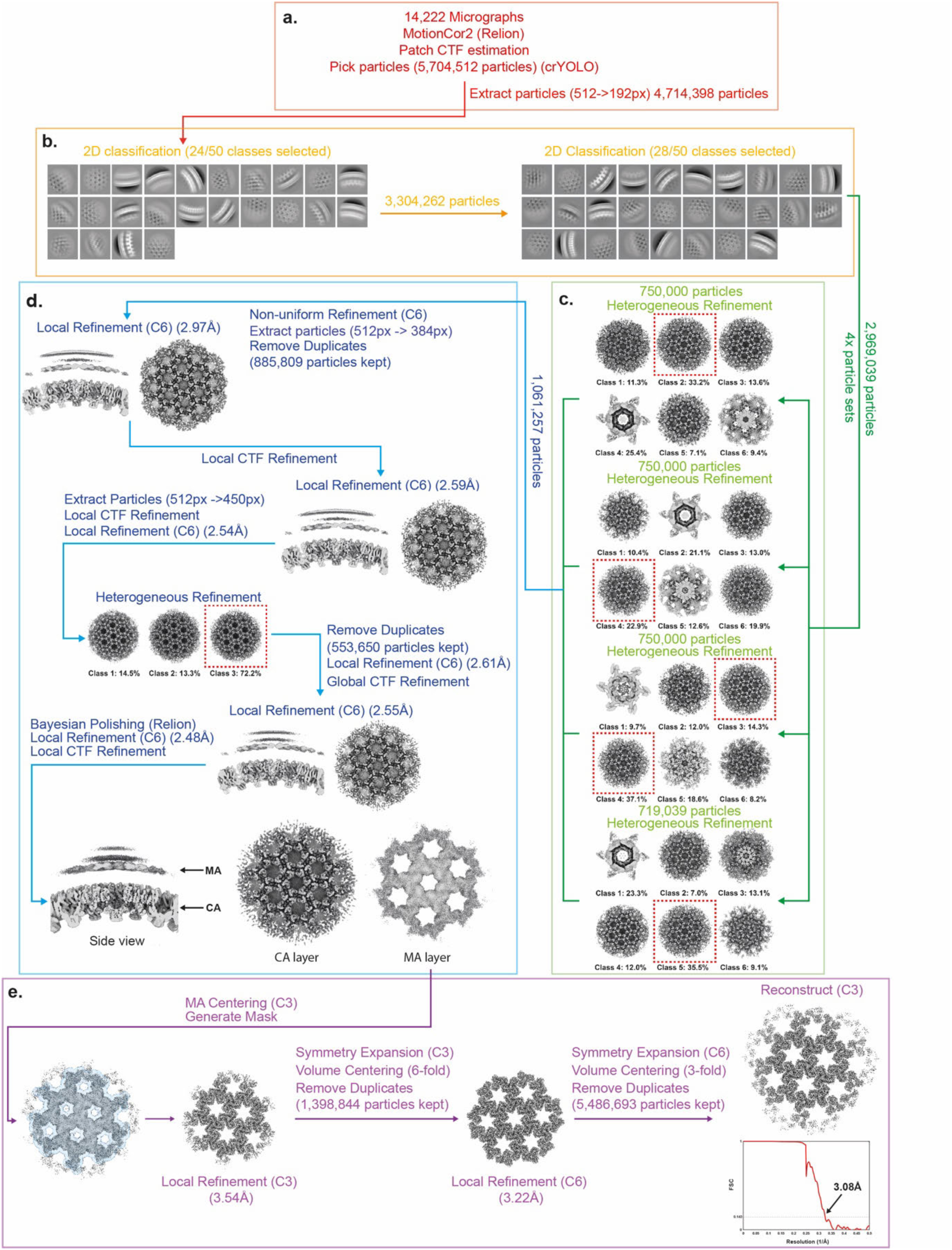
Single-particle cryo-EM processing for MA-SP1 MA lattice. For a detailed description of the steps illustrated see Materials and Methods **a,** EM micrograph preprocessing, particle picking and particle extraction. **b,** 2D classification of the Gag layer. **c,** Heterogeneous refinement of the CA layer. **d,** 3D alignment and refinement of the CA layer. **e,** 3D alignment and refinement of the MA layer, and FSC.

**Extended Data Fig. 5:**
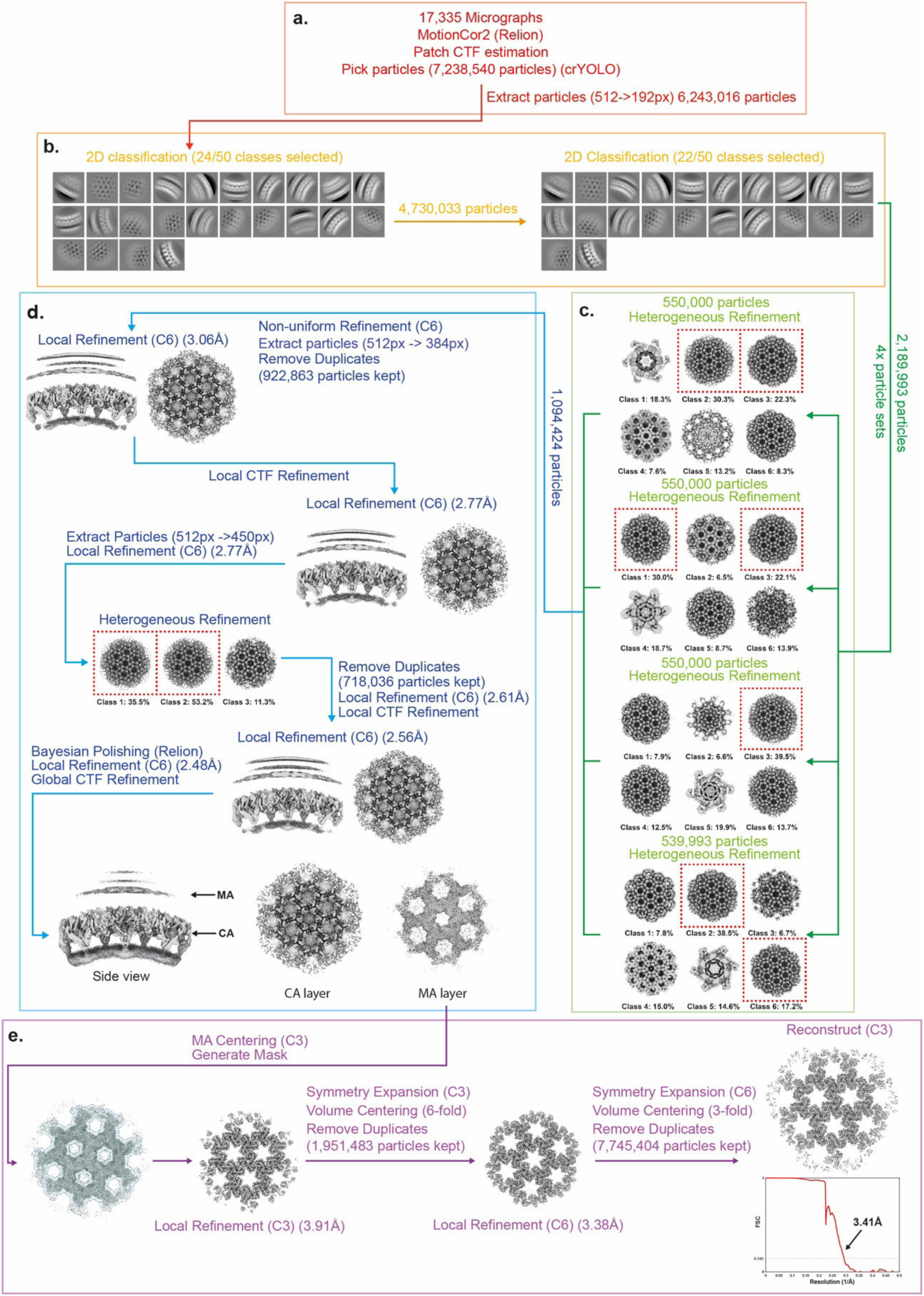
Single-particle cryo-EM processing for MA-NC MA lattice. The overall processing strategy of the MA-NC MA lattice was the same as that of the MA-SP1, except for the details specified in the figure. **a,** EM micrograph preprocessing, particle picking and particle extraction. **b,** 2D classification of the Gag layer. **c,** Heterogeneous refinement of the CA layer. **d,** 3D alignment and refinement of the CA layer. **e,** 3D alignment and refinement of the MA layer, and FSC.

**Extended Data Fig. 6:**
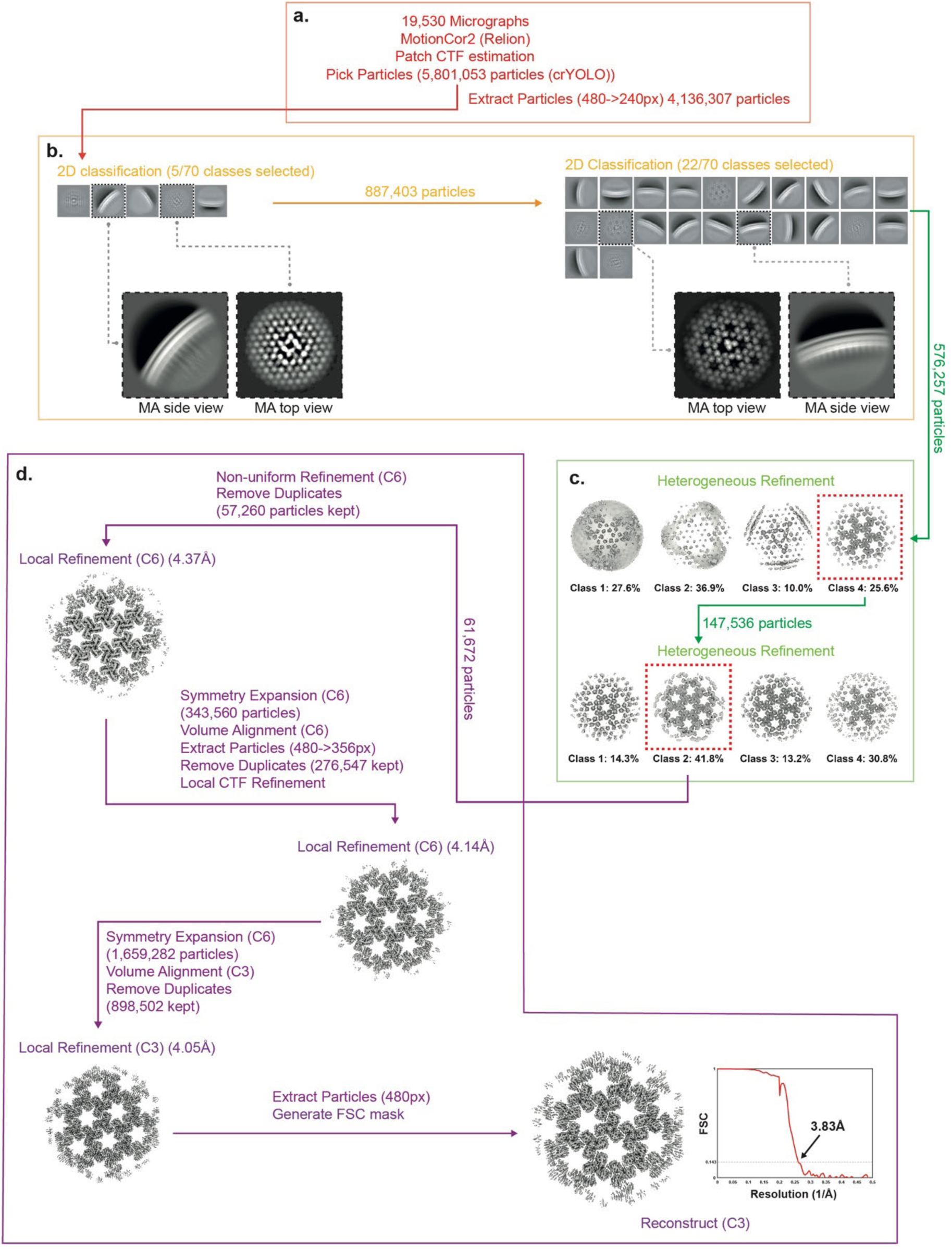
Single-particle cryo-EM processing for Mature (WT) MA lattice. For a detailed description of the steps illustrated see Materials and Methods **a,** EM micrograph preprocessing, particle picking and particle extraction. **b,** 2D classification of the MA layer. **c,** Heterogeneous refinement of the MA layer. **d,** 3D alignment and refinement of the MA layer, and FSC.

**Extended Data Fig. 7:**
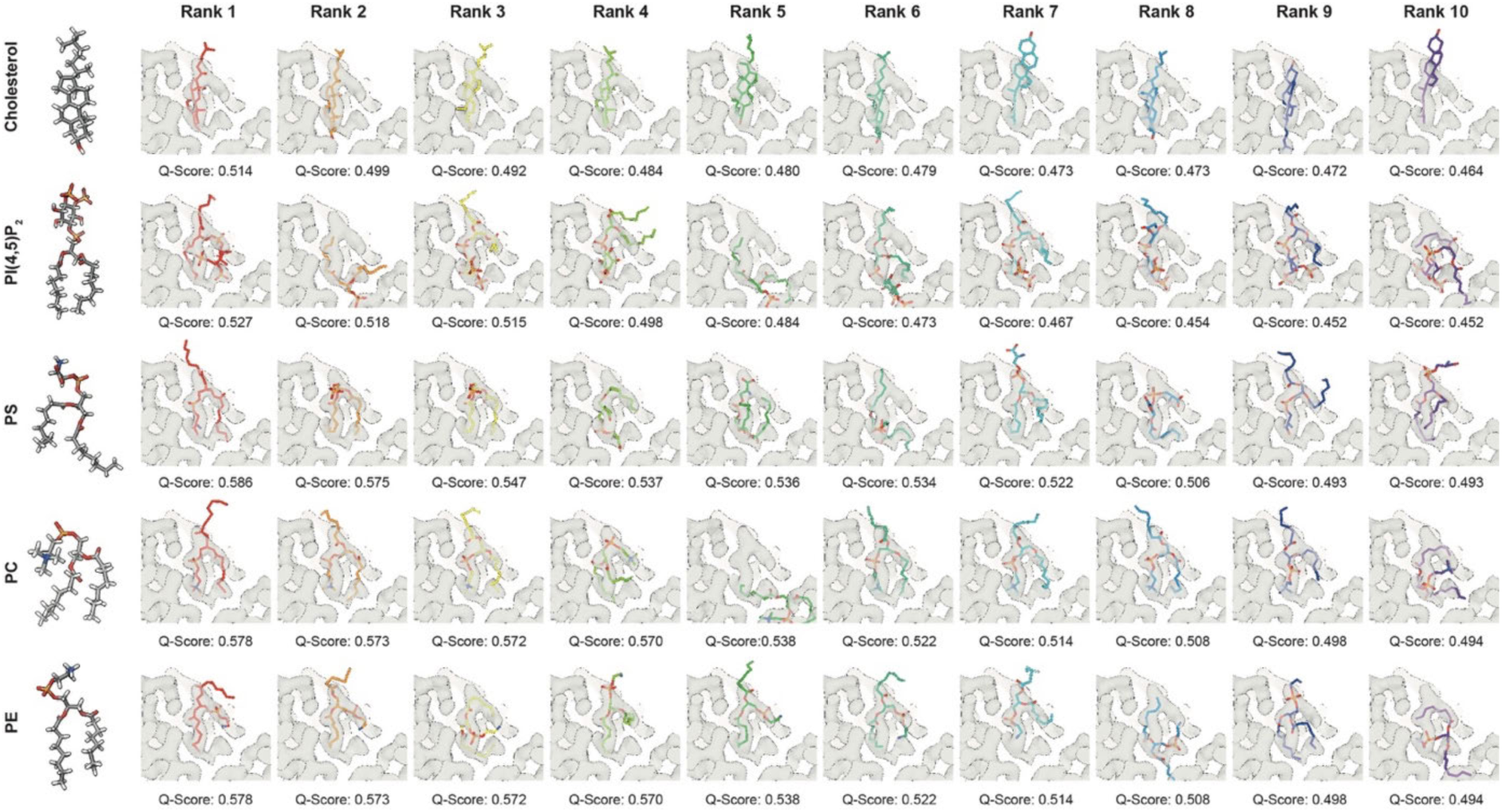
Further lipids do not fit into the side pocket density. Automated flexible ligand fitting was performed into the MA-SP1 ligand density using RosettaEmerald software ^53^. The ligands tested were Cholesterol, PI(4,5)P_2_, Phosphatidyl Serine (PS), Phosphatidyl Choline (PC) and Phosphatidyl Ethanolamine (PE). These ligands were chosen as they constitute the only lipids present in the HIV-1 viral membrane that are abundant enough to potentially completely occupy all MA binding sites (∼2000), as according to Mücksch et al., 2019 ^65^. The 10 resulting highest scoring fits (Q-Score) are shown in order of their rank. We identified no fit that could adequately account for the observed side pocket density.

**Extended Data Fig. 8:**
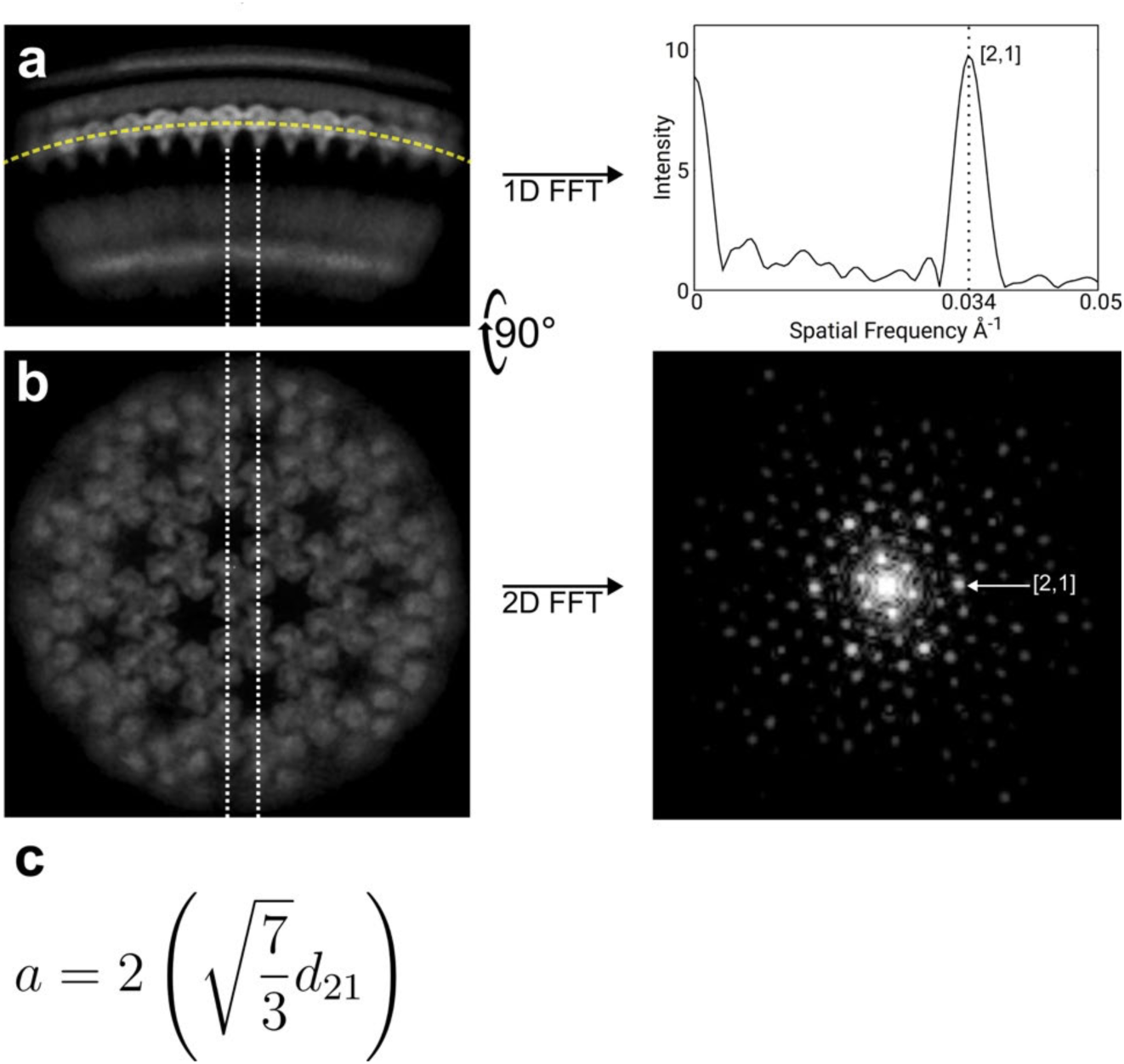
Further details of 2D class Fourier analysis of HIV-1 cleavage mutants. **a,** Side-view simulated projection of the mature matrix map of the HIV-1 MA-SP1 cleavage mutant. The orientation of the map is close to that observed in the 2D class averages used in the 1D Fourier analysis of matrix lattice. 1D fast Fourier transform (FFT) of pixel values extracted along the matrix layer produces a single peak at a spatial frequency of 0.034 Å^-1^ which corresponds to a mature lattice. **b,** Top-view (90-degree rotation of a. along the x-axis) simulated projection of the map. The matrix molecules oriented along the dotted lines produce strong reflection at h,k [2,1] as visible in the 2D FFT of the simulated projection. This is the same [2,1] reflection observed in the 1D FFT of the matrix side-view in a. **c,** Equation defining the relationship between the unit cell parameter *a* and the interplanar distance *d* of reflection h,k = [2,1].

**Extended Data Fig. 9:**
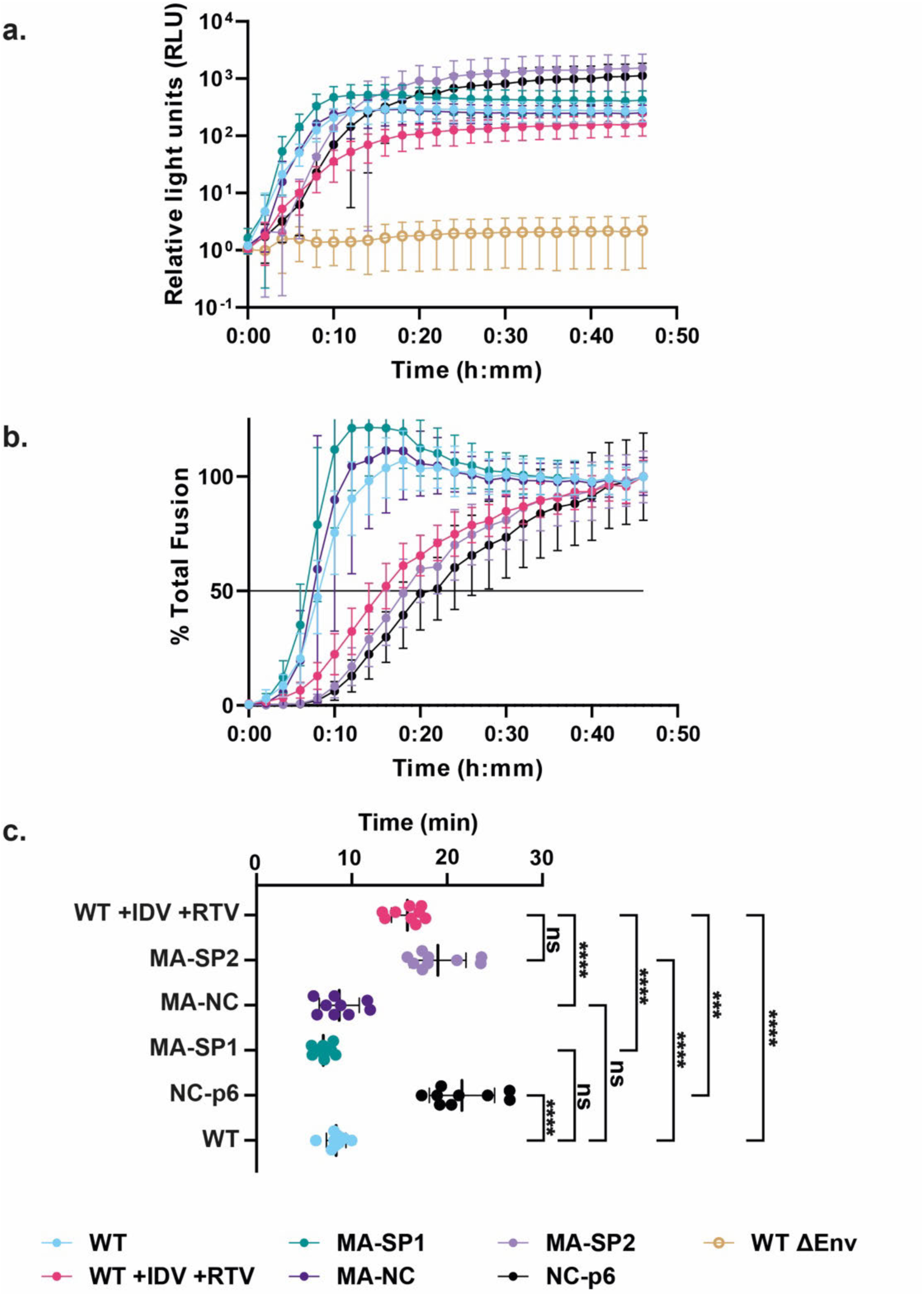
SP2 release is required for WT-like fusion kinetics. **a,** Graph of total luciferase signal measured in relative light units for indicated WT Gag virus-like-particles (VLPs), protease inhibitor-treated VLPs, and Gag cleavage mutant VLPs, all with WT JRFL Env. Readings were measured every 2 min from 0 min to 46 min. Blue = WT Gag VLPs, red = WT Gag VLPs produced in the presence of the protease inhibitors Indinavir (IDV) and Ritonavir (RTV), green = MA-SP1 Gag VLPs, dark purple = MA-NC Gag VLPs, lilac = MA-SP2 Gag VLPs, black = NC-p6 Gag VLPs, yellow open circle = WT Gag ΔEnv VLPs. Plotted points are the means from 3 biological replicates with 3 technical replicates per biological replicate for an n = 9. Each biological replicate represents one aliquot from a bulk preparation of viral particles. Error bars = s.d. **b,** Graph of percent of total fusion for each sample. Each sample is normalized with its own luciferase signal at 46 min as 100%. Samples are labelled as in (**a**). Black line represents 50% of total fusion observed. **c,** Histogram of fusion T_1/2_ for indicated VLPs produced in the presence of protease inhibitors Indinavir (IDV) and Ritonavir (RTV), Gag cleavage mutant VLPs, and WT Gag VLPs, all with WT JRFL Env. Mean ± s.d.: 15.83 ± 1.671 (WT +IDV +RTV), 19.04 ± 2.943 (MA-SP2), 8.677 ± 2.101 (MA-NC), 7.024 ± 0.9382 (MA-SP1), 21.56 ± 3.418 (NC-p6), 8.357 ± 1.022 (WT). Panel (**c**) is reproduced from Fig. 2d, but coloured in accordance with (**b**) and (**c**).

**Extended Data Fig. 10:**
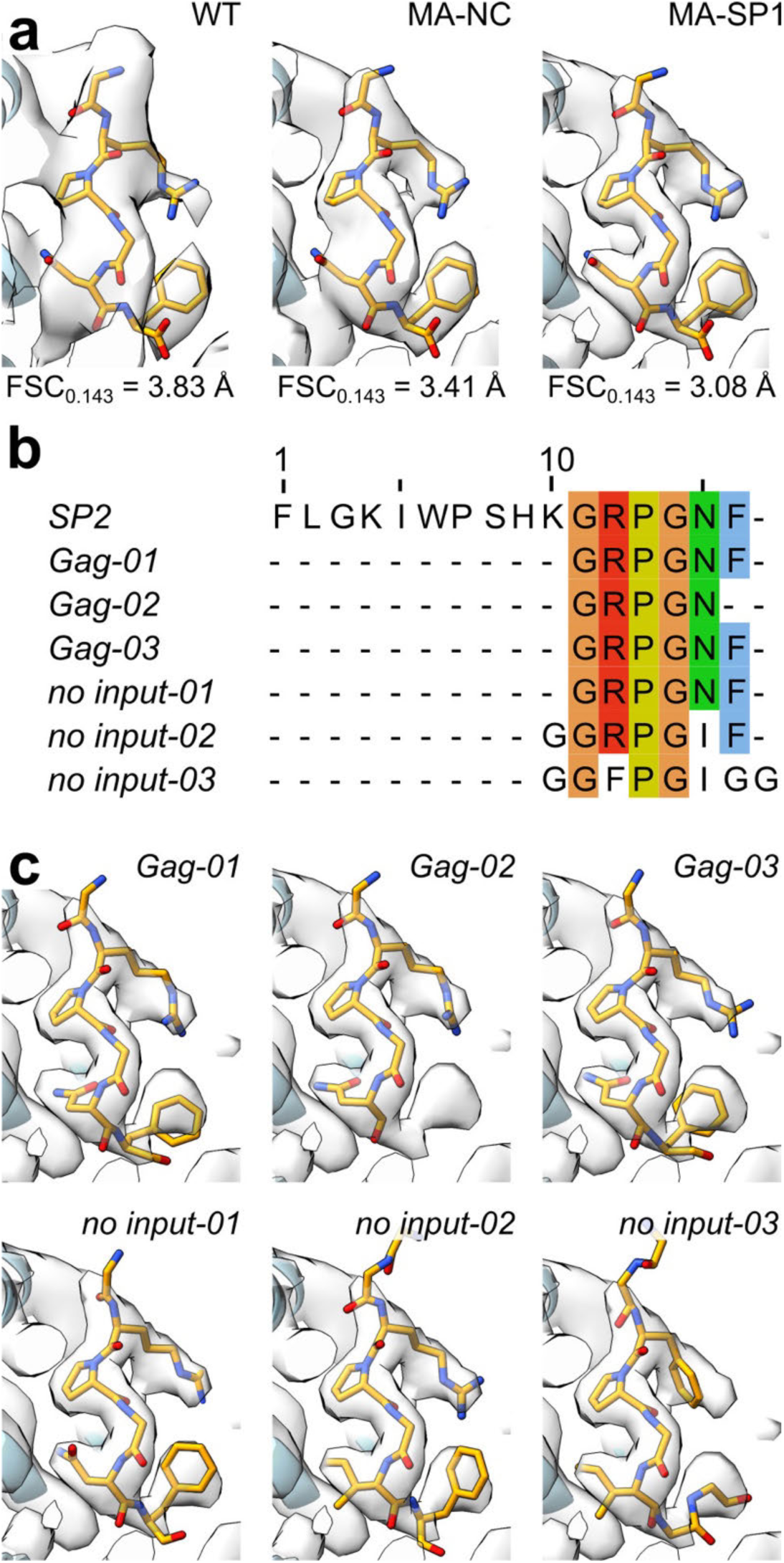
Automated model building of the density corresponding to SP2 using ModelAngelo. **a,** Atomic models of SP2 C-terminal residues fit into the side pocket densities of all determined mature MA lattice reconstructions (WT, MA-NC and MA-SP1). **b**, Multiple sequence alignment of automatically predicted sequences in the side pocket MA density by ModelAngelo ^59^. The first sequence is that of a full-length SP2 and the sequences below are either those predicted by ModelAngelo with the HIV-1 Gag sequence as input or with no sequence input. **c,** Atomic models built by ModelAngelo into the side pocket densities of the central MA trimer corresponding to the sequence predictions in a.

**Extended Data Fig. 11:**
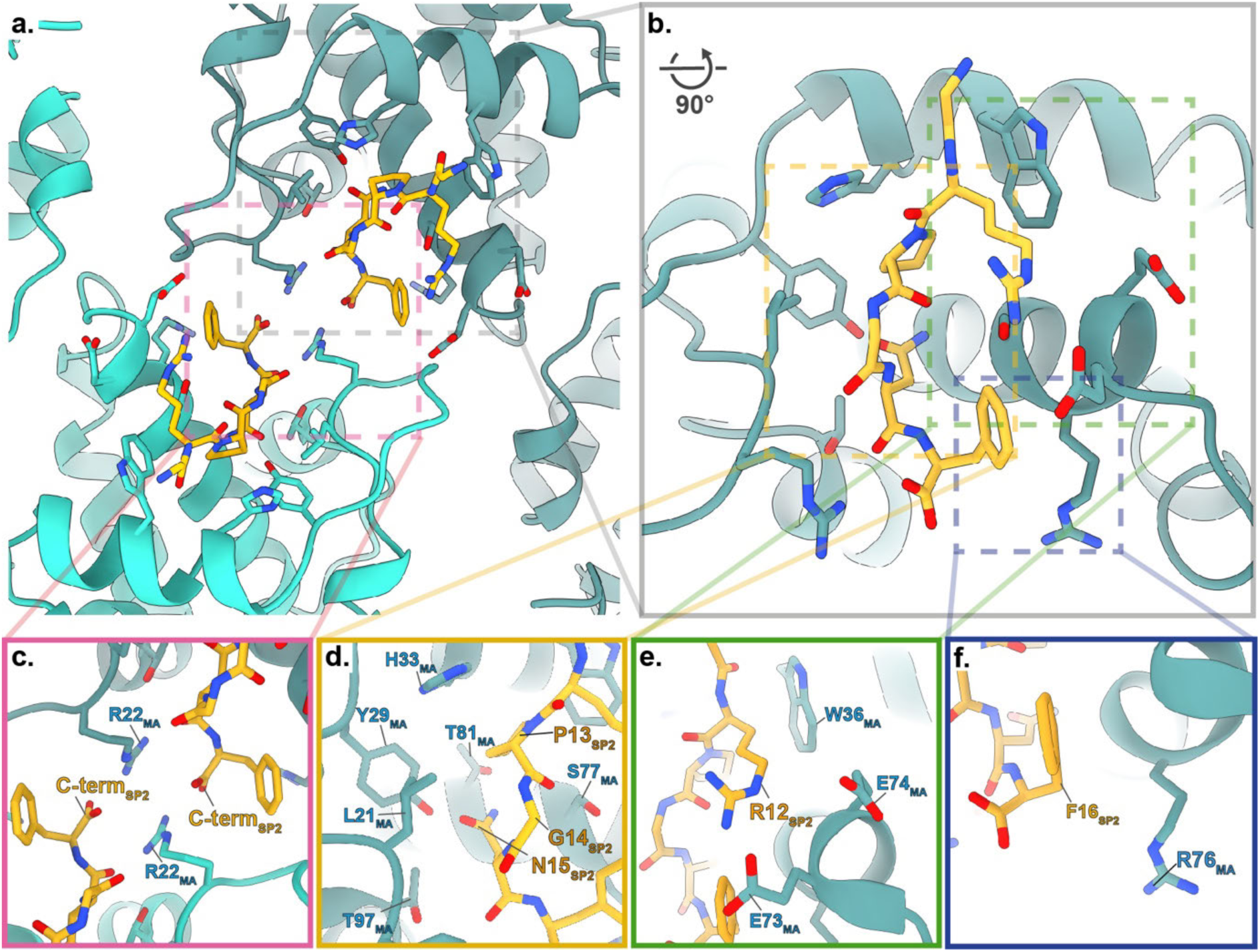
Structural details of the interaction between MA and SP2. **a,** Atomic model of the dimeric interface of the mature matrix lattice, with bound SP2, viewed from outside the virus (MA; Blue, SP2; orange). **b,** Rotated and zoomed in view of SP2 within the MA binding pocket from the side. **c,** Zoomed in view of A. The CTD of two adjacent SP2 chains and R22 from the two opposing MA molecules form a favourable electrostatic interaction network across the symmetry axis. This interaction likely results in the favouring of the mature MA lattice configuration upon SP2 binding to MA. **d-f,** Zoomed in views of b. **d,** Central SP2 binding motif residues P13, G14 and N15 sit in a hydrophobic cleft in the MA side pocket, formed by residues in helices 2, 4 and 5 as well as in the helix_1-2_ loop. **e,** Binding of SP2 R12 is supported by a hydrophobic contact with W36 and an electrostatic interaction with residues E73 and E74. **f,** Binding of SP2 F16 is facilitated by a hydrophobic contact with R76.

**Extended Data Fig. 12:**
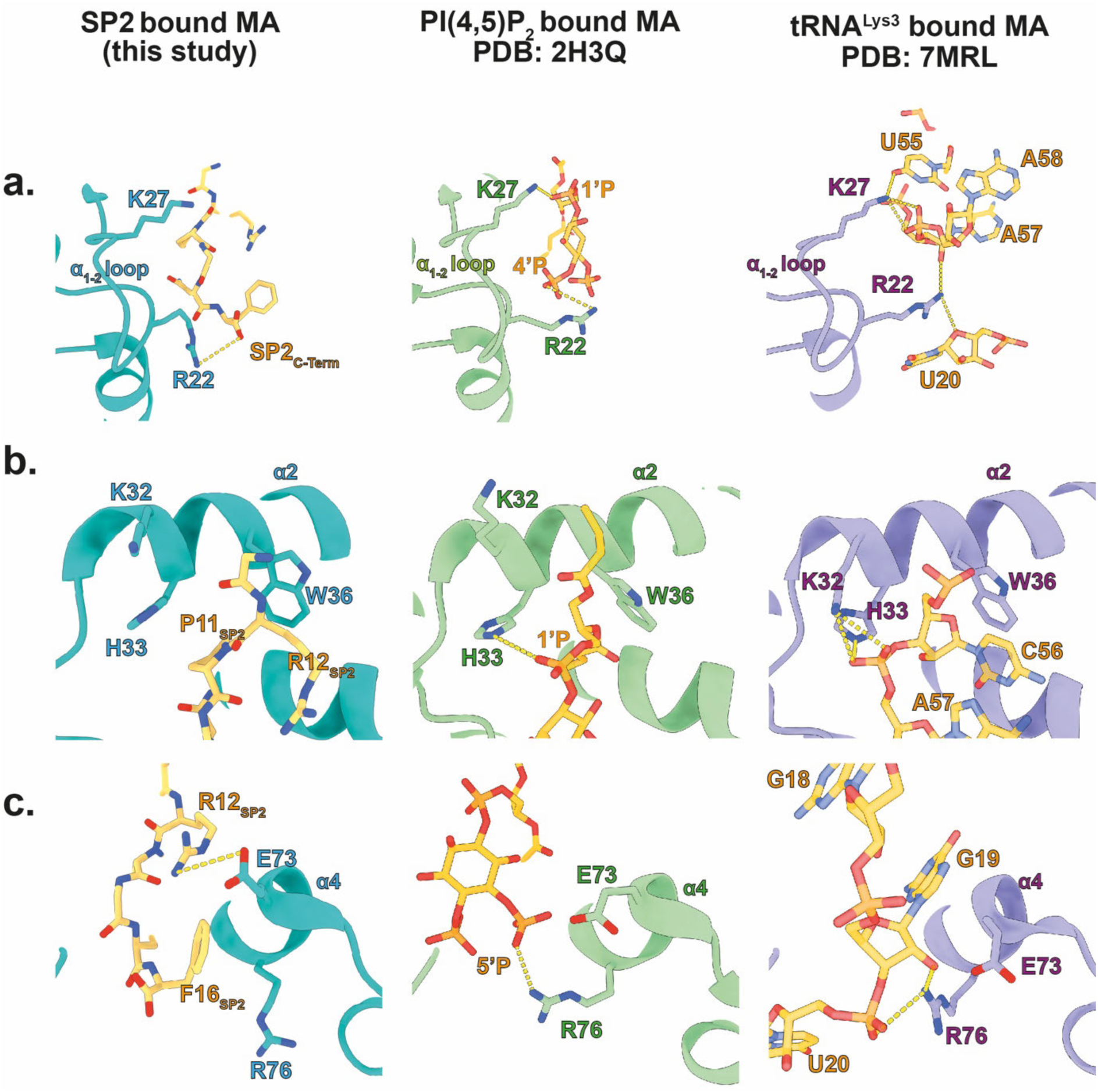
Comparison of SP2, PI(4,5)P_2_ and tRNA^Lys3^ binding to MA. **a,** MA bound to SP2 (left. MA: pink, SP2: orange): R22 in the helix_1-2_ loop forms an electrostatic interaction with the C-terminal carboxyl group of SP2, whereas K27 does not interact with SP2 (left). MA bound to PI(4,5)P_2_ ^11^ (middle. MA green; PI(4,5)P_2_ orange): R22 and K27 interact with the PI(4,5)P_2_ 4’-phosphate and bridging 1’-phosphodiester respectively. MA bound to tRNA (right. MA purple; tRNA^Lys3^ orange): R22 forms an interaction with a 2’-OH ribose moiety (tA57) and a 4O ribose position on the tRNA backbone (tU20) whereas K27 interacts with a backbone phosphate group (tA58) (right) ^23^. **b,** Left: Helix_2_ residues, H33 and W36, form π-stacking interactions with P11 and the R12 backbone amide of SP2 respectively. K32 does not interact with SP2. Middle: H33 forms an electrostatic interaction with the 1’ phosphate of PI(4,5)P_2_. Right: Both K32 and H33 form electrostatic interactions with backbone phosphate of the tRNA (tA57), whereas W36 forms a π stacking interaction with tC56. **c,** Left: Helix_4_ residue E73 forms a salt bridge with R12 of SP2. F16 of SP2 packs against the acyl tail of R76. Middle: R76 forms an electrostatic interaction with the 5’-phosphate of PI(4,5)P_2_. Right: R76 interacts with a backbone phosphate of tRNA as well as a ribose 2’-OH (tG16).

**Extended Data Fig. 13:**
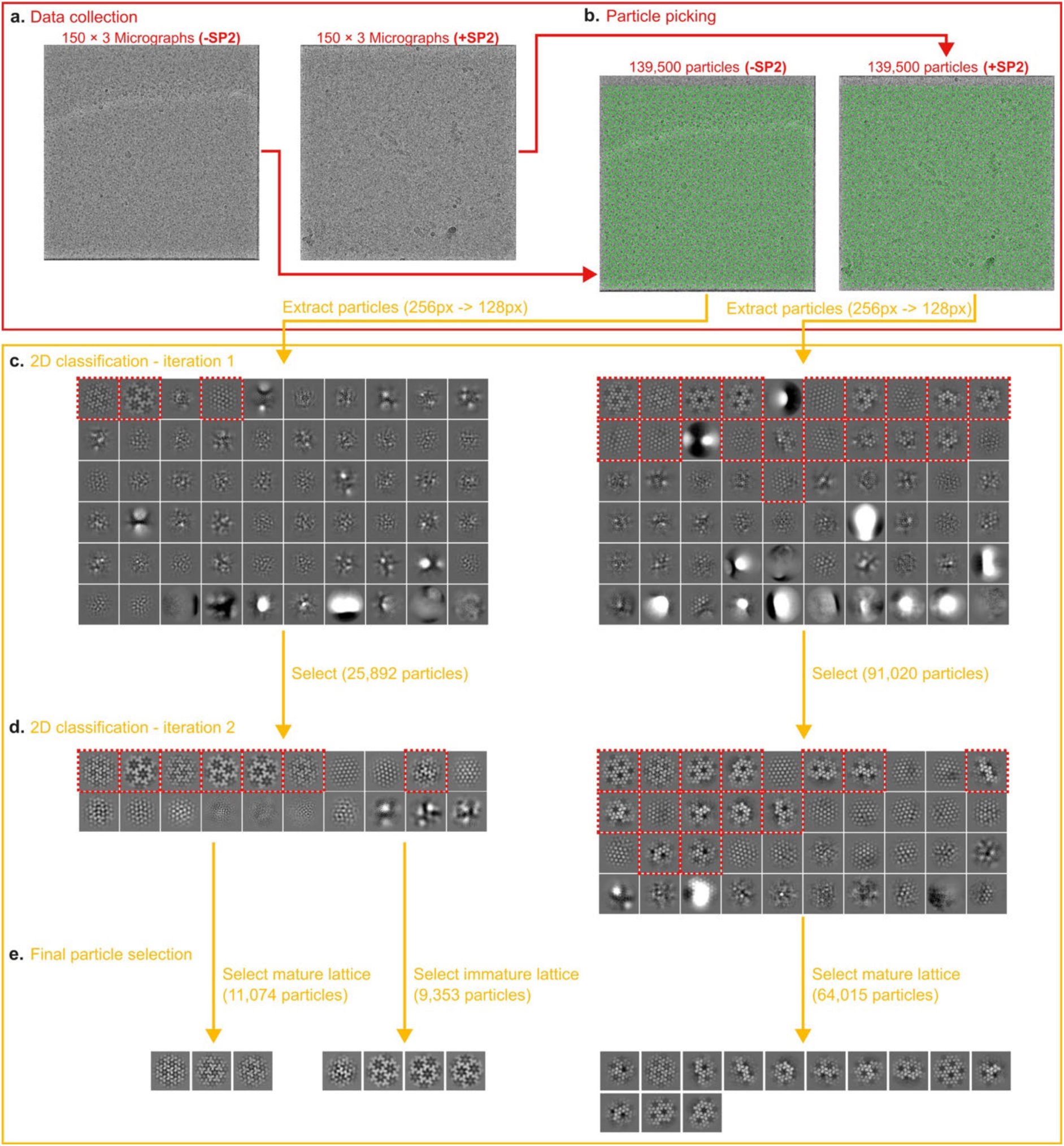
Data processing scheme of the 2D crystallography experiment. **a,** 150 micrographs were collected randomly from each grid. Six grids were prepared simultaneously, three grids of the myrMA -SP2 control group and three grids of the myrMA +SP2 experimental group. **b,** Particles were picked randomly as a grid of points and extracted in cryosparc. **c,** The extracted particles were subjected to an initial round of 2D classification. 2D classes showing any kind of MA lattice were selected and subjected to a second round of 2D classification. **d,** Classes with mature and immature lattices were selected and particles belonging to these classes were used in the final statistical analysis.

**Extended Data Fig. 14:**
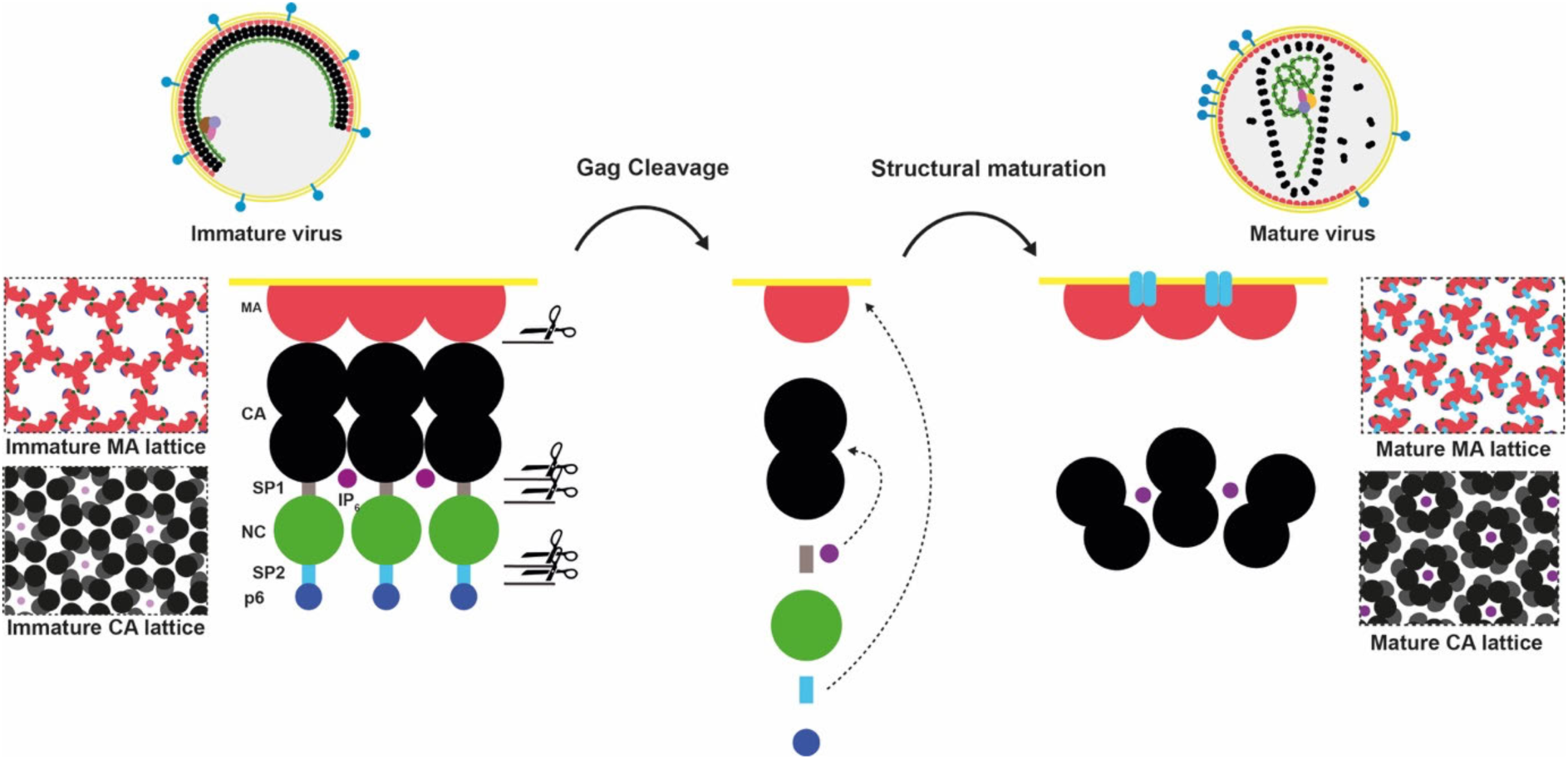
Schematic illustration of the mechanism of MA and CA maturation. In the immature virus, MA and CA both form ‘immature’ hexameric lattices. Upon viral maturation, which is initiated by the activation of the viral protease (PR), Gag is cleaved into its constituent domains; MA, CA, SP1, NC, SP2 and p6. In the case of MA, subsequent structural transition to the mature lattice is induced upon binding of cleaved SP2 to MA. In the case of CA, cleavage of SP1 leads to the release of IP_6_ which binds free CA to promote the formation of the mature CA lattice.

**Extended Data Table. 1:** Summary table of Cryo-EM data collection and processing statistics. * Additional particles were derived during processing by lattice expansion (see Materials and Methods)

|  | HIV-1<br>Immature (PR-)<br>EMD:XXXXX | HIV-1<br>MA-SP1<br>EMD:51769 | HIV-1<br>Mature (WT)<br>EMD:XXXXX | HIV-1<br>MA-NC<br>EMD:XXXXX |
| --- | --- | --- | --- | --- |
| **Data collection and processing** |  |  |  |  |
| Magnification | 130,000x | 130,000x | 130,000x | 130,000x |
| Voltage (kV) | 300 | 300 | 300 | 300 |
| Electron exposure (e<sup>-</sup>/Å<sup>2</sup>) | 40 | 40 | 40 | 40 |
| Defocus range (μm) | -0.6 to -2.6 | -0.6 to -2.6 | -0.6 to -2.6 | -0.6 to -2.6 |
| Pixel size (Å) | 0.95 | 0.95 | 0.95 | 0.95 |
| Movies (no.) | 9,942 | 14,222 | 19,530 | 17,335 |
| Initial particle images (no.) | 3,568,755 | 5,704,512 | 5,801,053 | 7,238,540 |
| Symmetry imposed | C3 | C3 | C3 | C3 |
| Final particle images (no.)* | 174,245 | 5,486,693 | 889,805 | 7,745,404 |
| Map resolution (Å) | 5.83 | 3.08 | 3.83 | 3.41 |
| FSC threshold | 0.143 | 0.143 | 0.143 | 0.143 |

**Extended Data Table. 2:**
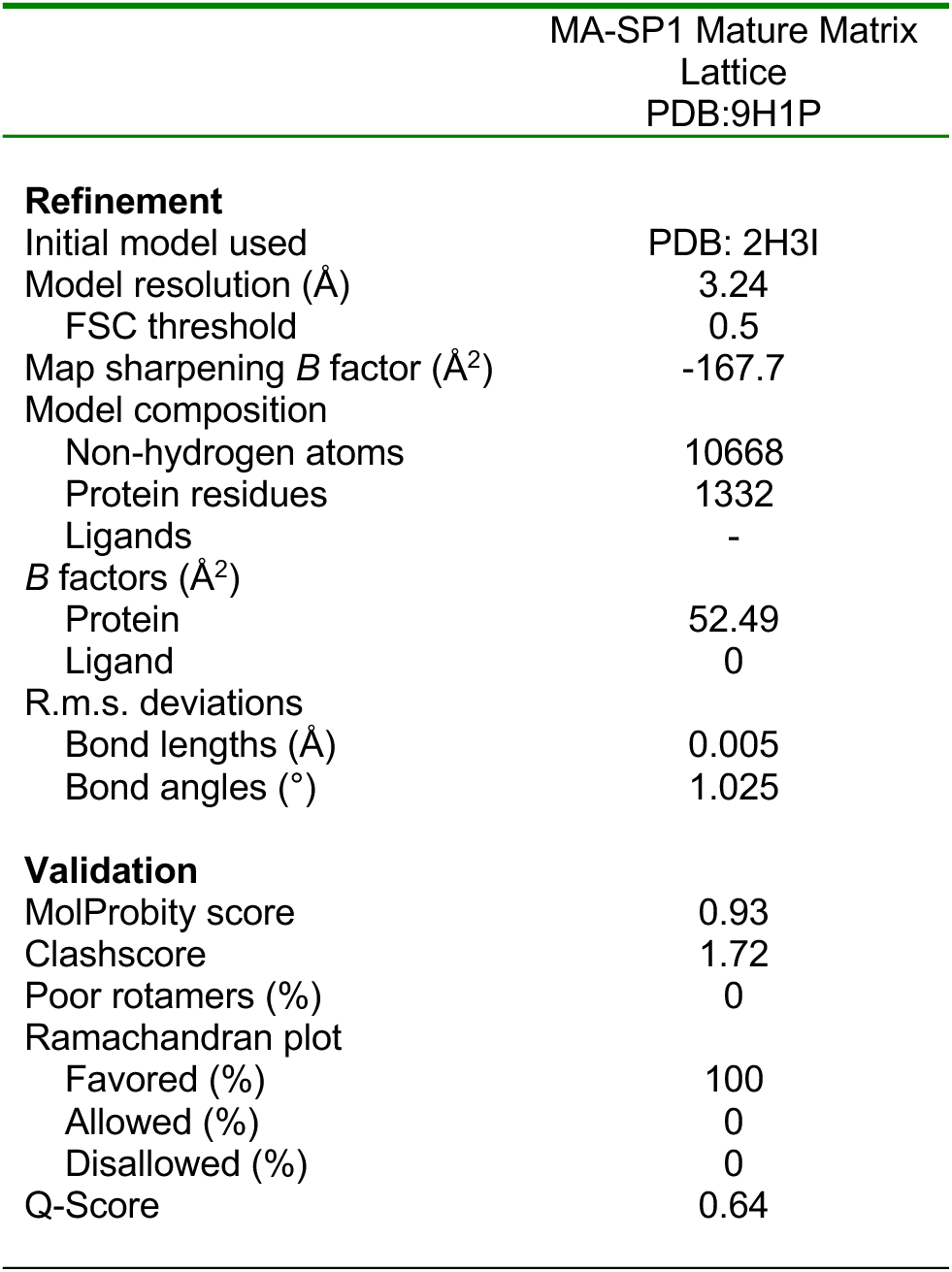
Summary table of model building and refinement statics of mature matrix lattice in complex with SP2 (HIV-1 MA-SP1).

